# Translational control through differential ribosome pausing during amino acid limitation in mammalian cells

**DOI:** 10.1101/321448

**Authors:** Alicia M. Darnell, Arvind R. Subramaniam, Erin K. O’Shea

## Abstract

Limitation for amino acids is thought to regulate translation in mammalian cells primarily by signaling through the kinases mTORC1 and GCN2. We find that limitation for the amino acid arginine causes a selective loss of tRNA charging, which regulates translation through ribosome pausing at two of six arginine codons. Surprisingly, limitation for leucine, an essential and abundant amino acid in protein, results in little or no ribosome pausing. Chemical and genetic perturbation of mTORC1 and GCN2 signaling revealed that their robust response to leucine limitation prevents ribosome pausing, while an insufficient response to arginine limitation led to loss of arginine tRNA charging and ribosome pausing. Codon-specific ribosome pausing decreased protein production and triggered premature ribosome termination without significantly reducing mRNA levels. Together, our results suggest that amino acids which are not optimally sensed by the mTORC1 and GCN2 pathways still regulate translation through an evolutionarily conserved mechanism based on codon-specific ribosome pausing.

## Introduction

Cells need to regulate anabolic processes to maintain homeostasis in the face of fluctuating nutrient levels. Of these processes, protein synthesis consumes the highest fraction of nutrients and energy stores in proliferating cells (Buttgereit and Brand, 1995; Hosios et al., 2016), and is therefore tightly controlled in response to fluctuations in the levels of its amino acid substrates. In eukaryotic cells, amino acid limitation is sensed by two evolutionarily conserved signaling pathways anchored around the kinases mechanistic Target Of Rapamycin in Complex 1 (mTORC1) (Saxton and Sabatini, 2017) and General Control Nonderepressible 2 (GCN2) (Berlanga et al., 1999; Hinnebusch and Natarajan, 2002). Amino acid limitation inhibits mTORC1 signaling (Hara et al., 1998) and activates GCN2 signaling (Sood et al., 2000), which reduces overall protein synthesis rate through a decrease in the rate of ribosome initiation on mRNA transcripts (Sonenberg and Hinnebusch, 2009). The failure of either pathway to respond to amino acid limitation can lead to cell death, particularly in nutrient-challenged contexts such as tumors (Nofal et al., 2017; Ye et al., 2010) or neonates (Efeyan et al., 2013; Zhang et al., 2002), underscoring the importance of their regulatory control over protein synthesis in maintaining cellular homeostasis.

The mTORC1 and GCN2 pathways both respond strongly to simultaneous limitation for all 20 amino acids (Kimball, 2002), yet their responses to fluctuations in the levels of individual amino acids are markedly different. mTORC1 signaling is highly sensitive to fluctuations in leucine levels, and to a lesser extent, to arginine and glutamine levels (Hara et al., 1998). By contrast, GCN2 kinase, which senses amino acid limitation by binding uncharged tRNAs, has a similar affinity for different tRNAs (Dong et al., 2000; Zaborske et al., 2010), but variation in its response to limitation for individual amino acids is nonetheless detected in activation of the downstream transcriptional program (Jousse et al., 2000; Tang et al., 2015). How these variegated mTORC1 and GCN2 responses are integrated, and whether they are sufficient, to regulate protein synthesis rate during individual amino acid limitation is poorly understood. This is of increasing importance as there is growing evidence that many cancers exhibit dependence on single amino acids for growth or metastasis (Hattori et al., 2017; Jain et al., 2012; Knott et al., 2018; Krall et al., 2016; Loayza-Puch et al., 2016a; Possemato et al., 2011; Scott et al., 2000; Wise and Thompson, 2010).

In addition to mTORC1- and GCN2-mediated regulation of translation initiation, amino acid limitation can affect protein synthesis by reducing the elongation rate of ribosomes. In bacteria, limitation for single auxotrophic amino acids causes selective loss of tRNA isoacceptor charging and thus ribosome pausing at a subset of synonymous codons cognate to the limiting amino acid (Dittmar et al., 2005; Subramaniam et al., 2013a). This ribosome pausing results in abortive termination and a consequent decrease in protein expression (Ferrin and Subramaniam, 2017; Subramaniam et al., 2013b, 2014). Notably, the codons at which ribosomes pause during amino acid limitation are not necessarily rare codons or decoded by low abundance tRNA isoacceptors (Subramaniam et al., 2013b, 2013a, 2014). Ribosomes pause during histidine limitation in yeast, but whether this pausing is codon-specific, and its impact on protein expression, are not known (Guydosh and Green, 2014). Ribosome pausing has also been observed in pathological mammalian states, including in a mouse model of neurodegeneration (Ishimura et al., 2014), and in patient-derived cancer tissues (Loayza-Puch et al., 2016a). However, the factors that drive ribosome pausing in these cancer cells are unclear and difficult to parse *in vitro*. Further, the codon-specificity and effect of ribosome pausing on protein expression have not been studied in mammalian systems, though codon usage frequency and tRNA levels have been implicated in the regulation of ribosome elongation rate and protein production during metastasis, differentiation, and amino acid limitation (Gingold et al., 2014; Goodarzi et al., 2016; Saikia et al., 2016). However, ribosome profiling studies have failed to find evidence for a simple relationship between codon usage, tRNA levels and ribosome density in mammalian cells (Ingolia et al., 2011; Qian et al., 2012).

Here, we investigated how amino acid signaling pathways and codon usage interact to regulate protein synthesis in response to limitation for single amino acids across multiple human cell lines. We focused on limitation for two amino acids, leucine and arginine, which can both regulate protein synthesis by acting as direct signals to the mTORC1 complex (Chantranupong et al., 2016; Wolfson et al., 2016). Upon arginine limitation, we found that a stereotypical pattern of ribosome pausing emerges at the same two out of six synonymous arginine codons across cell lines, suggesting that arginine becomes a rate-limiting substrate in protein synthesis. Intriguingly, there was little to no ribosome slow-down at any of the six leucine codons upon limitation for leucine, even though it is an essential amino acid. The hierarchy of ribosome pausing at synonymous arginine codons was not correlated with codon usage or genomic tRNA copy number, but followed the selective loss of arginine isoacceptor tRNA charging. By perturbing amino acid signaling, we established that tRNA charging loss and ribosome pausing are driven by an inadequate response to amino acid limitation through the mTORC1 and GCN2 pathways. We found that codon-specific ribosome pausing decreases both the rate of global protein synthesis as well as protein expression from individual mRNAs. Further, severe pausing caused by loss of the mTORC1 and GCN2 signaling responses to amino acid limitation triggers the premature termination of protein synthesis.

Our study provides a mechanistic dissection of the cause and consequences of ribosome pausing due to amino acid limitation in mammalian cells. We reveal an evolutionarily conserved role for synonymous codon-specific ribosome pausing in the regulation of protein synthesis during amino acid limitation, a phenomenon which has been previously observed only in bacteria (Subramaniam et al., 2013b, 2014). However, we discovered a layer of complexity in this process that is unique to mammalian cells – quantitative differences in the activity of amino acid signaling pathways result in qualitative differences in ribosome pausing upon limitation for the two amino acids arginine and leucine. By establishing a molecular framework relating amino acid depletion, tRNA charging, ribosome elongation, and protein expression, our work provides a rational starting point from which to dissect the cellular phenotype of disease states, such as cancers, that experience nutrient limitation and exhibit dysregulated ribosome dynamics (Ishimura et al., 2014; Loayza-Puch et al., 2016b).

## Results

### 1.1. Ribosomes pause at specific synonymous codons upon limitation for arginine but not leucine

To systematically explore the effect of individual amino acid depletion on translation in mammalian cells, we performed ribosome profiling (Ingolia et al., 2009, 2012) in three human cell lines – HEK293T, HeLa and HCT116 – during limitation for either leucine or arginine. Although ribonuclease I (RNaseI) is typically used to generate monosome-bound RNA footprints for ribosome profiling (Ingolia et al., 2012), we found that micrococcal nuclease (MNase) treatment better preserved monosome integrity (Supp. Fig. 1A-C, Methods), and sequencing the resulting footprints (Supp. Fig. 1D) produced ribosome profiling libraries with reads enriched in coding regions and displaying three nucleotide periodicity, despite a broader read length distribution, as previously reported (Dunn et al., 2013; Reid et al., 2015) (Supp. Fig. 1E-G). After sequencing, we quantified the net increase in normalized average ribosome footprint density in the window around each of the 61 sense codons as a measure of the change in elongation kinetics of ribosomes upon amino acid limitation (Fig. 1A; Supp. Fig. 1H, Methods).

**Fig. 1.**
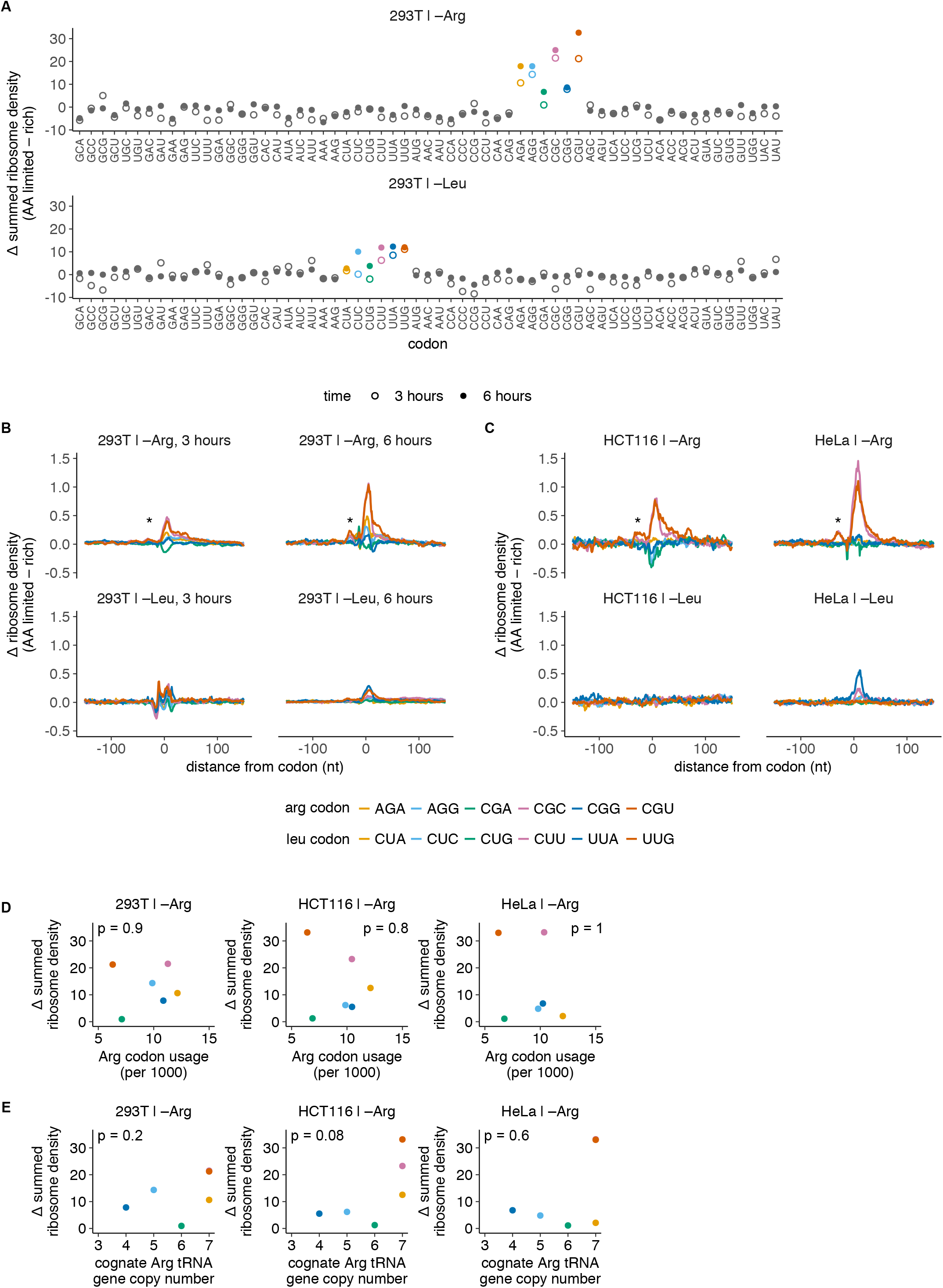
Codon-specific ribosome pausing during limitation for arginine, but not leucine. (**A-C**) Changes in codon-specific ribosome density for HEK293T cells, HCT116, and HeLa cells following 3 or 6 hours of arginine or leucine limitation, measured using ribosome profiling. Ribosome density for each codon is calculated relative to the mean footprint density for each coding sequence, and is averaged over all occurrences of each of the 61 sense codons across detectably expressed transcripts. The difference in ribosome density between amino acid limited and rich conditions across a 150 nt window around each codon is either summed (A) or shown as such (B,C). Asterisk indicates satellite peaks reflecting collision of the trailing ribosome with the paused ribosome. (**D**) Usage frequency of Arg codons in the HEK293T, HCT116, and HeLa transcriptomes following 3 hours of arginine limitation (as shown in Supp. Fig. 1J) is compared to the summed change in ribosome density upon arginine limitation (as shown in A for HEK293T, and Supp. Fig. 1H for HCT116, HeLa). p indicates p-value of Spearman’s rank coefficient, ρ (HEK293T; ρ = –0.1, p = 0.9. HCT116; ρ = –0.1, p = 0.8. HeLa; ρ = 0.03, p = 1). (**E**) Genomic copy number of the cognate tRNA for each Arg codon (Chan and Lowe, 2016) (as shown in Supp. Fig. 1K) compared to the change in ribosome density upon arginine limitation (as shown in A for HEK293T, and Supp. Fig. 1H for HCT116, HeLa) (HEK293T; ρ = 0.58, p = 0.2. HCT116; ρ = 0.76, p = 0.08. HeLa; ρ = 0.27, p = 0.6).

Upon arginine limitation for three hours, two of the six arginine codons–CGC and CGU–had a substantial increase in ribosome density across all three cell lines (Fig. 1A,C; Supp. Fig. 1H). Ribosome pausing at these codons increased with prolonged amino acid limitation for six hours (Fig. 1B). None of the codons encoding the other 19 amino acids had increased ribosome density upon arginine limitation (Fig. 1A, Supp. Fig. 1H). Notably, we also observed smaller peaks in ribosome density approximately one ribosome footprint length (∼ 30 nucleotides) behind the major peaks at CGC and CGU codons (Fig. 1B,C; asterisks). Similar satellite peaks, presumably caused by collision of the trailing ribosome with the paused ribosome, have been previously observed during limitation for single amino acids in *E. coli* (Subramaniam et al., 2014) and in *S. cerevisiae* (Guydosh and Green, 2014).

In contrast, none of the six leucine codons displayed a consistent increase in ribosome density across all three cell lines in response to leucine limitation (Fig. 1A-C; Supp. Fig. 1H). Since leucine cannot be synthesized in these cells, we were surprised to find that ribosome elongation at leucine codons is largely unperturbed by leucine limitation. We considered the possibility that cells do not experience major changes in intracellular leucine levels upon its external limitation. However, direct measurement of cellular amino acid levels indicated that arginine and leucine levels fell close to the detection limit when they were each removed from the growth medium, suggesting that cells are effectively starved for both leucine and arginine in these conditions (Supp. Fig. 1I).

We then tested whether the selective increase in ribosome density upon arginine limitation correlated with simple measures of codon optimality or tRNA abundance, as hypothesized previously (Gingold et al., 2014; Kirchner and Ignatova, 2015; Saikia et al., 2016). The pausing hierarchy did not correlate significantly in any cell line with either transcriptomic codon usage (Fig. 1D, Supp. Fig. 1J,L) or genomic copy number of the cognate tRNA (Fig. 1E, Supp. Fig. 1K,M,N) (Kanaya et al., 1999) (Spearman’s rank correlation p-values displayed on plots). Nevertheless, the consistent hierarchy of codon-specific ribosome pausing upon arginine limitation, and its absence during leucine limitation, suggests a common principle underlying the emergence of ribosome pausing.

### 1.2. Cognate tRNA charging loss upon amino acid limitation sets the hierarchy of ribosome pausing at synonymous codons

As ribosome elongation rate at a codon depends on recruitment of the cognate charged tRNA, we expected that the arginine tRNA which decodes the two pause-inducing codons CGC and CGU, with the anticodon ACG (tRNA^Arg^_ACG_), would exhibit a greater charging loss upon arginine limitation than the isoacceptor arginine tRNAs that decode the remaining four arginine codons. In line with this expectation, tRNA^Arg^_ACG_ lost 70% of its charging upon arginine limitation in HEK293T cells (Fig. 2A, Supp. Fig. 2A). By contrast, tRNA^Arg^_CCG_ and tRNA^Arg^_UCG_, which decode the arginine codons CGG and CGA at which we did not observe strong pausing, lost less than 45% of their charging (Fig. 2A, Supp. Fig. 2A). All leucine tRNAs tested lost less than 40% of their charging upon leucine limitation, consistent with the observation that there is no ribosome pausing at leucine codons (Fig. 2B, Supp. Fig. 2B). As expected, arginine and leucine tRNAs were between 75% to 90% charged during growth in rich conditions, and upon limitation for a non-cognate amino acid (Fig. 2A,B). Charging loss was also more severe for tRNA^Arg^_ACG_ than a leucine tRNA in the HCT116 cell line (Supp. Fig. 2C). Overall we found a positive correlation between the change in ribosome density at a codon and the loss in charging of its cognate tRNA upon limitation for an amino acid (Spearman’s rank correlation coefficient *ρ* = 0.7, p = 0.015; Fig. 2C). Our results suggest that ribosomes begin to pause at a codon only when a majority of the cognate charged tRNA is depleted.

**Fig. 2.**
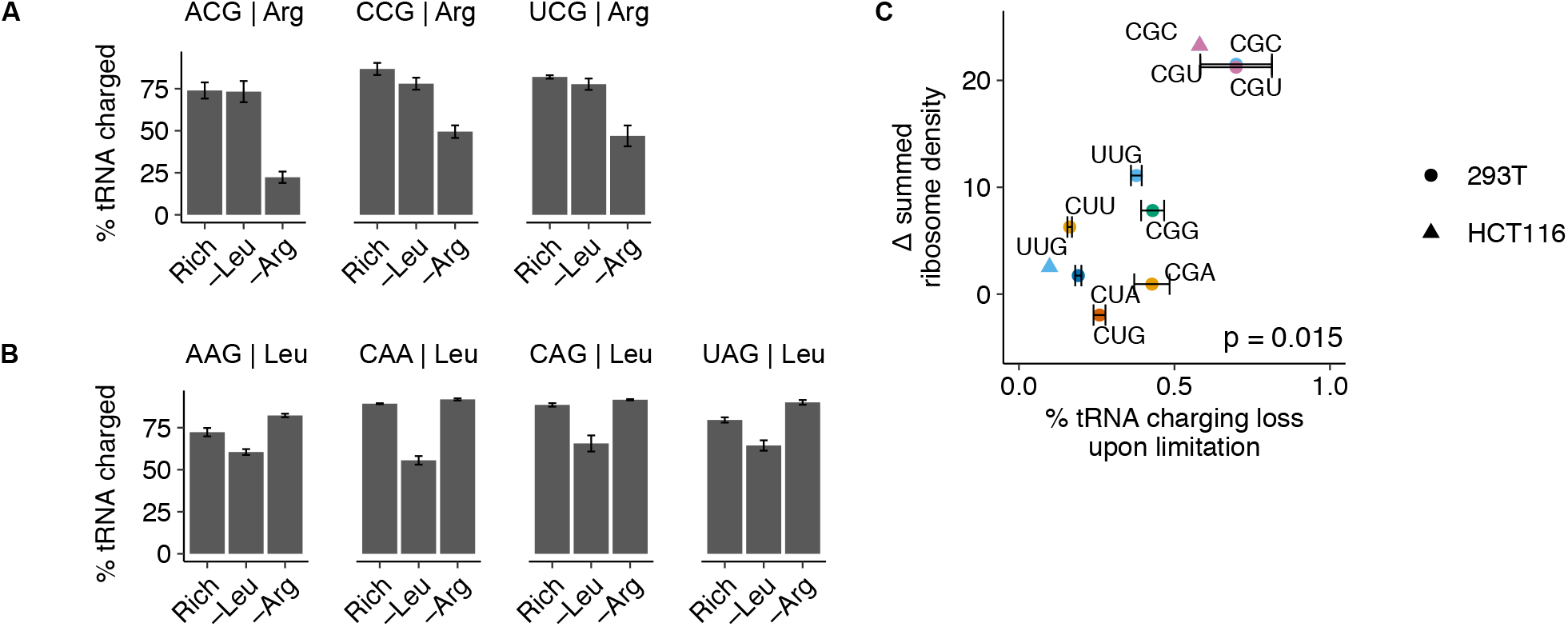
Selective loss of cognate tRNA charging during arginine limitation. (**A,B**) tRNA charging levels for 3 Arg tRNAs (A) and 4 Leu tRNAs (B) in HEK293T cells following 3 hours of leucine or arginine limitation or growth in rich medium (calculated as described in Supp. Fig. 2A). tRNA anticodon and isotype are indicated above plots; error bars represent the standard error of the mean from three technical replicate experiments (see Supp. Fig. 2A,B for representative northern blots and Supp. Fig. 1M for codon-tRNA pairs). (**C**) Summed change in ribosome density at arginine and leucine codons for HEK293T cells (Fig. 1A) plotted against the loss in charging for cognate tRNA (for those measured) following arginine or leucine limitation, respectively. p indicates p-value of Spearman’s rank coefficient, ρ (ρ = 0.87, p = 0.005, N = 8).

### 1.3. The mTORC1 and GCN2 pathways respond divergently to arginine limitation

We next examined whether the loss of charged tRNA and emergence of ribosome pausing during arginine but not during leucine limitation might be related to the amino acid signaling response through the GCN2 and mTORC1 pathways, given that these pathways are presumed to sense amino acid levels and co-ordinately regulate protein synthesis in order to maintain intracellular amino acid homeostasis (Bröer and Bröer, 2017). Consistent with previous reports (Hara et al., 1998), we observed greater mTORC1 inhibition during limitation for leucine in comparison to arginine – levels of the mTORC1 target phosphorylated S6 kinase 1 (P∼S6K) fell by 75% during leucine limitation, but only 45% during arginine limitation in HEK293T cells (Fig. 3A). Levels of the S6K target phosphorylated ribosomal protein S6 (P∼RPS6) correspondingly reflected this differential mTORC1 response (Supp. Fig. 3A,B). GCN2 signaling was strongly activated during limitation for both amino acids in these cells – levels of the GCN2 target phosphorylated eIF2α (P∼eIF2α) increased to a similar extent (Fig. 3B).

**Fig. 3.**
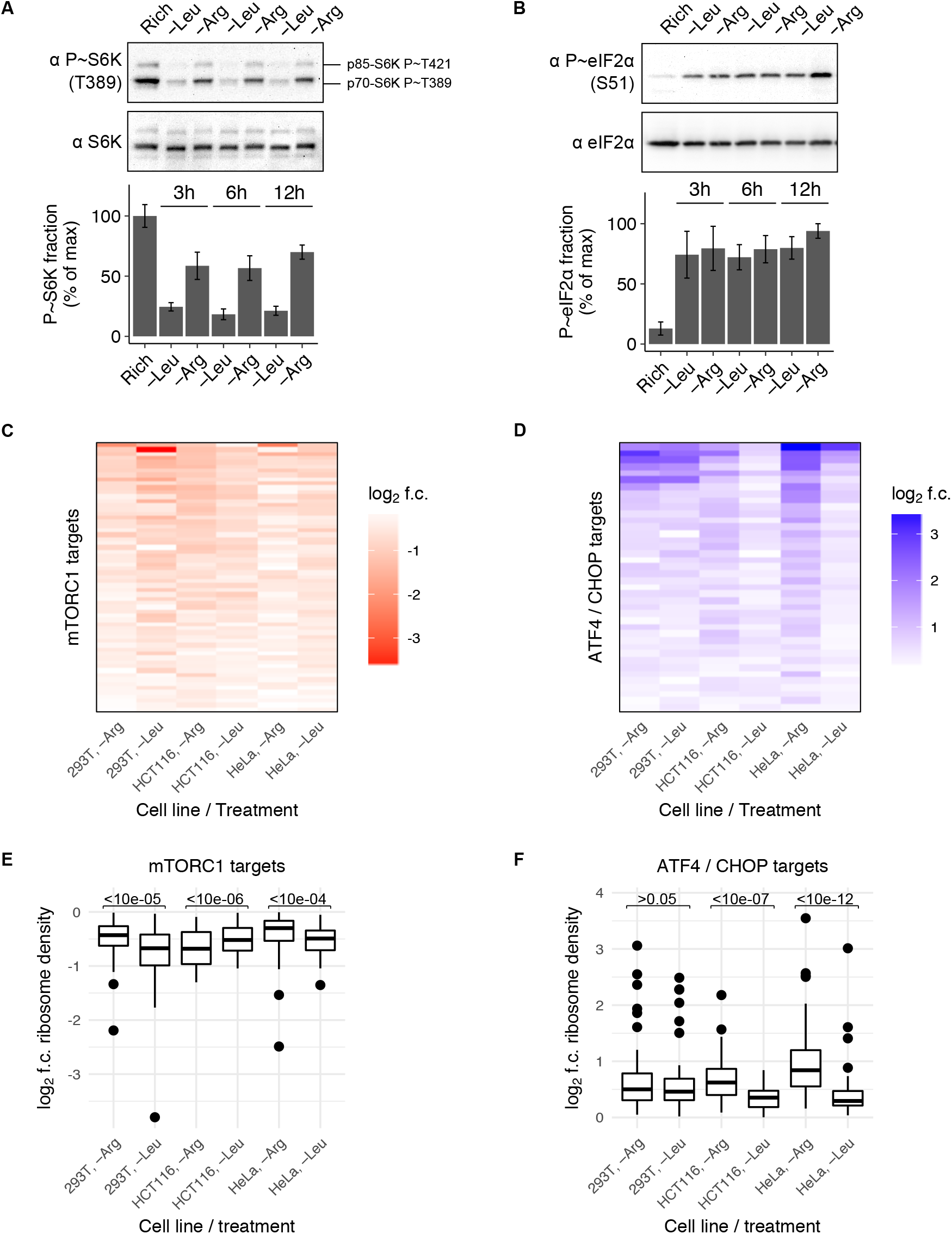
Divergent response of mTORC1 and GCN2 signaling pathways to arginine versus leucine limitation. (**A,B**) Representative western blots for phosphorylated and total levels of the mTORC1 target p70 S6 kinase 1 (S6K) (**A**) or the GCN2 target eIF2α (**B**) in HEK293T cells after growth in rich medium or after 3, 6, or 12 hours of leucine or arginine limitation. Bar graph shows the fraction of protein that is phosphorylated in each condition, relative to the maximum; this normalized phosphorylation index was found first for each sample on one blot and then averaged between blots from replicate experiments. Error bars represent the standard error of the mean from three technical replicate experiments. (**C,D**) Heatmap of log_2_ fold-change (f.c.) in ribosome density for mRNA targets of translational downregulation due to mTORC1 inhibition (Hsieh et al., 2012) (C) or of transcriptional or translational upregulation due to GCN2 activation (Han et al., 2013) (D), following 3 hours of arginine or leucine limitation relative to growth in rich medium for HEK293T, HCT116, and HeLa cells. Only targets with a log_2_ fold change of < 0, for mTORC1, or > 0, for ATF4/CHOP (transcription factor effectors downstream of GCN2 activation), upon amino acid limitation in all conditions and cell lines were considered. In HEK293T cells, 46/63 (73%), in HCT116 cells, 14/63 (22%), and in HeLa cells, 45/63 (71%) of mTORC1 targets had higher ribosome density upon arginine than leucine cells, 40/40 (100%) of GCN2 targets had higher ribosome density upon arginine than leucine limitation. (**E,F**) Box plot of the log_2_ fold change for each mTORC1 (E) or ATF4/CHOP (F) target upon amino acid limitation (as shown in C,D). A two-sided Wilcoxon signed rank test with continuity correction was performed to test the null hypothesis that the median difference (μ) in the log_2_ fold change for each target between arginine and leucine limitation was equal to zero. The resulting p-value is shown above each comparison and indicates whether there is a significant difference in the signaling response to arginine versus leucine limitation. In HEK293T, HCT116, and HeLa cells, the mTORC1 signaling response was 1.2-, 0.9-, and 1.1-fold higher during limitation for arginine, respectively (E) and the GCN2 signaling response was 1-, 1,2, and 1.5-fold higher during limitation for arginine, respectively (F).

Kinase activity in HEK293T cells mirrored downstream changes in ribosome density on mRNA targets of the mTORC1 pathway. 46 of 63 mRNAs that are translationally repressed by mTORC1 inhibition (Hsieh et al., 2012; Thoreen et al., 2012) had lower ribosome density during limitation for leucine than arginine (Fig. 3C,E; Supp. Fig. 3C,E,G; Wilcoxon signed rank test p = 1.2e-05). Similarly, mTORC1 signaling was more repressed during limitation for leucine in HeLa cells (Fig. 3C,E; Supp. Fig. 3G; Wilcoxon signed rank test p = 0.0003). This pattern was reversed in HCT116 cells, in which there was little mTORC1 or GCN2 signaling response to leucine limitation (Fig. 3C-F; Supp. Fig. 3G), consistent with our observation that leucine tRNA charging is largely unaffected by leucine limitation (Supp. Fig. 2C).

Comparing downstream changes in ribosome density on mRNA targets of ATF4 and CHOP, transcriptional effectors downstream of GCN2 (Han et al., 2013), during arginine versus leucine limitation revealed subtle but consistent differential activation of GCN2. In HEK293T cells, GCN2 signaling was similarly activated during limitation for leucine and arginine; 26 out of 40 of mRNA targets of ATF4 and CHOP, were more upregulated upon limitation for arginine than leucine (Fig. 3D,F; Supp. Fig. 3D,F,G; Wilcoxon signed rank test p = 0.33). However, GCN2 became significantly more activated during arginine limitation after a longer duration of amino acid limitation (Supp. Fig. 3D,F; Wilcoxon signed rank test p = 5.7e-4), which also increased ribosome pausing (Fig. 1A,B). In addition, GCN2 was more activated during limitation for arginine in the HCT116 and HeLa cell lines (Wilcoxon signed rank test p = 9.3e-07 and 1.8e-12, respectively) (Fig. 3D,F; Supp. Fig. 3G). GCN2 was generally most responsive in the conditions and cell lines in which ribosome pausing was most severe, consistent with the recent observation that GCN2 may be activated downstream of ribosome pausing (Ishimura et al., 2016).

Overall, the variability of the signaling responses across all three cell lines was surprising, given that we observed a conserved signature of ribosome pausing. However, if pausing is determined by the extent to which the amino acid supply and demand are matched under each condition, it may be the totality of the signaling response, rather than the activity of each single pathway, that regulates this balance. We sought to test this idea in the HEK293T cell line, in which ribosome pausing emerges only during arginine limitation, in the context of a relatively weaker overall signaling response than leucine limitation.

### 1.4. An insufficient mTORC1 and GCN2 response to amino acid limitation induces ribosome pausing

The mTORC1 and GCN2 pathways inhibit the initiation phase of protein synthesis in response to amino acid limitation (Ma and Blenis, 2009; Sonenberg and Hinnebusch, 2009). Reducing initiation rate should also lower the number of elongating ribosomes – a major source of demand for the cytosolic amino acid pool – thereby determining the consumption rate of a limiting amino acid. If the strength of their combined signaling response is too weak to sufficiently reduce arginine consumption during its limitation, tRNA charging loss and ribosome pausing could result. Specifically, if residual mTORC1 activity and/or inadequate activation of GCN2 drives amino acid consumption and thus loss of tRNA charging and ribosome pausing, we hypothesized that increasing the response of these pathways would reduce pausing upon arginine limitation, and conversely, that decreasing their response would induce pausing upon leucine limitation. To test this hypothesis, we employed chemical and genetic methods to perturb the mTORC1 and GCN2 responses to arginine and leucine limitation in HEK293T cells, and determined the resulting changes to tRNA charging and ribosome pausing.

We first inhibited mTORC1 kinase activity using the catalytic site inhibitor Torin1 (Liu et al., 2010; Thoreen et al., 2009) (Fig. 4A) during both arginine and leucine limitation, and found that charging of all arginine and leucine tRNAs tested increased back to baseline rich conditions levels (Supp. Fig. 4A). Torin1 treatment also prevented an increase in ribosome density at any codon upon arginine or leucine limitation (Fig. 4B, Supp. Fig. 4B), demonstrating that mTORC1 inhibition during amino acid limitation is sufficient to block depletion of the cognate charged tRNA fraction and ribosome pausing.

**Fig. 4.**
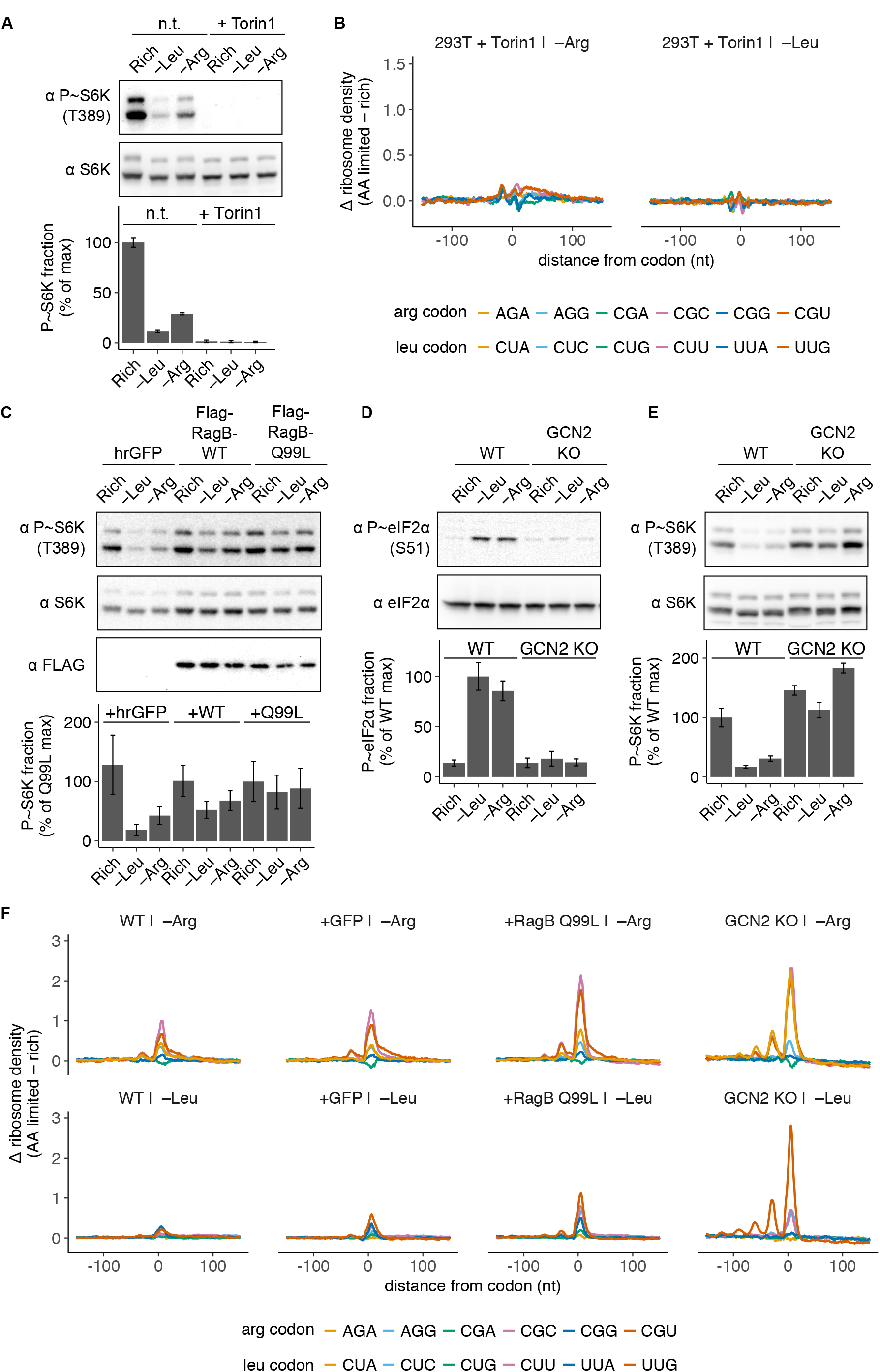
Signaling through the mTORC1 and GCN2 pathways regulates the magnitude of ribosome pausing during amino acid limitation. (**A**) Representative western blots for phosphorylated and total S6K in HEK293T cells after growth in rich medium or limitation for leucine or arginine for 3 hours, in the presence or absence of 250 nM Torin1. Bar graph shows fraction of protein that is phosphorylated, relative to rich medium; error bars represent the standard error of the mean from three technical replicate experiments. (**B**) Changes in codon-specific ribosome density in HEK293T cells expressing a fluorescent reporter protein (hrGFP, as shown in C) after 3 hours of arginine or leucine limitation with 250 nM Torin1, relative to the maximum. (**C**) Representative western blots for phosphorylated S6K, total S6K, and Flag epitope after growth in rich medium, or 3 hours of leucine or arginine limitation in HEK293T cells stably expressing either hrGFP, Flag-RagB-WT, or Flag-RagB-Q99L. RagB-Q99L is a dominant positive mutant of RagB, an that is phosphorylated, relative to the maximum in the RagB-Q99L cell line; error bars represent the standard error of the mean from three technical replicate experiments. (**D,E**) Representative western blots for phosphorylated and total eIF2α (D) or S6K (E) after growth in rich medium, or 3 hours of leucine or arginine limitation in wild-type (WT) or GCN2 knock-out (KO) HEK293T cells. Bar graphs show fraction of protein that is phosphorylated, relative to the maximum in WT cells; error bars represent the standard error of the mean from three technical replicate experiments. (**F**) Changes in codon-specific ribosome density for WT HEK293T, hrGFP, Flag-RagB-Q99L, and GCN2 KO cell lines following 6 hours of limitation for arginine or leucine.

Next, we tested whether loss of the mTORC1 response to amino acid limitation would exacerbate tRNA charging loss and ribosome pausing. Towards this, we rendered mTORC1 kinase insensitive to amino acid levels by stable overexpression of a constitutively active form of its upstream regulator, RagB GTPase (RagB-Q99L) (Sancak et al., 2008) (Fig. 4C). The RagB-Q99L cell line exhibited reduced leucine tRNA charging during leucine limitation; charging fell to 22% for tRNA^Leu^_CAA_, which decodes the codon UUG (Supp. Fig. 4C). By comparing charging for this tRNA during leucine limitation in the RagB-Q99L cell line to a control line that over-expressed humanized *R. reniformis* fluorescent protein (hrGFP), we concluded that constitutive mTORC1 activity increased charging loss due to leucine limitation by 50%. Charging was also reduced 36% due to constitutive mTORC1 activity for tRNA^Leu^_AAG_, which decodes CUU (Supp. Fig. 4C). Concordantly, minor ribosome pausing was detected at the leucine codons UUG and CUU (Supp. Fig. 4D). However, little difference was detected in arginine tRNA charging or ribosome pausing at arginine codons upon arginine limitation (Supp. Fig. 4C,D), and we thus repeated these measurements after 6 hours, rather than 3 hours, of amino acid limitation to reveal any effects on translation that might become more pronounced over time.

After 6 hours of limitation for leucine, the RagB-Q99L cell line exhibited further reduced charging of leucine tRNAs compared to control cell lines; charging fell as low as 18% for tRNA^Leu^_CAA_ (Supp. Fig. 4E) and ribosome pausing emerged at the cognate leucine codon UUG as well as the CUC and CUU codons (Fig. 4F; Supp. Fig. 4F). Similarly, during arginine limitation, the proportion of charged tRNA^Arg^_ACG_ fell to 19% (Supp. Fig. 4E) and ribosome pausing increased at the cognate arginine codons CGC and CGU (Fig. 4F; Supp. Fig. 4F). Ribosome pausing was also increased slightly in the hrGFP control cell line (Fig. 4F, Supp. Fig. 4F), possibly due to the translational burden of transgene overexpression (Elf et al., 2003). In summary, constitutive mTORC1 activation in the RagB-Q99L cell line significantly worsened tRNA charging loss and exacerbated ribosome pausing during both leucine and arginine limitation.

We next investigated the role of GCN2 in ribosome pausing. We constructed a GCN2 knockout (GCN2 KO) cell line by CRISPR/Cas9 targeting (Cong et al., 2013) (Supp. Fig. 4G) in which the GCN2 kinase target eIF2α was not phosphorylated in response to amino acid limitation (Fig. 4D). GCN2 activation is necessary for inhibition of mTORC1 signaling upon leucine or arginine limitation in mouse as well as in murine and human cancer cell lines (Averous et al., 2016; Xiao et al., 2011), and we confirmed that there is no significant mTORC1 response to those conditions in our GCN2 KO cell line (Fig. 4E). tRNA charging loss and ribosome pausing were greatly amplified in the GCN2 KO cell line; tRNA^Leu^_CAA_ charging fell to only 14% upon leucine limitation (Supp. Fig. 4E) and ribosome density at the UUG leucine codon rose substantially, with a genome-wide average of 4 ribosomes stacked behind the paused ribosome (Fig. 4F, Supp. Fig. 4F). Pausing increased only slightly at the arginine CGC and CGU codons (Fig. 4F, Supp. Fig. 4F), although tRNA^Arg^_ACG_ charging continued to drop (Supp. Fig. 4E), indicating that pause duration is approaching an upper limit at these codons. Indeed, significant ribosome pausing emerged at the AGA arginine codon (Fig. 4F, top panel; Supp. Fig. 4F), suggesting that charging of a second arginine isoacceptor, tRNA^Arg^_UCU_, is exhausted upon arginine limitation in the GCN2 KO cell line. Together these results indicate that the absence of a response through the GCN2 or mTORC1 pathways during amino acid limitation is sufficient to deplete charged tRNA pools and induce extensive genome-wide ribosome pausing at cognate codons, consistent with our hypothesis that an insufficient signaling response to amino acid limitation can drive consumption of the limiting amino acid into a substrate-limiting regime for protein synthesis.

In addition to their control over translation, mTORC1 and GCN2 regulate other critical functions such as metabolism, autophagy, and cell division (Castilho et al., 2014; Laplante and Sabatini, 2013). In principle, regulation of these processes could affect intracellular amino acid levels. Hence, it is possible that mTORC1 and GCN2 determine whether ribosome pausing arises during amino acid limitation by controlling these processes in addition to, or instead of, by reducing translation. To test our hypothesis that arginine and leucine levels during their respective limitation are primarily determined by the demand from translation elongation, we briefly treated cells limited for arginine or leucine with the translation elongation inhibitor cycloheximide. This was sufficient to significantly restore tRNA^Leu^_CAA_ and tRNA^Arg^_ACG_ charging (Supp. Fig. 4H), indicating that the flux of arginine and leucine into translation is a key determinant of the cytosolic levels of these amino acids upon their limitation. Thus, the ribosome pausing outcome is likely determined by the translational control imposed downstream of mTORC1 and GCN2 during limitation for an amino acid.

### 1.5. Genome-wide ribosome pausing reduces global protein synthesis rate during arginine limitation

Having examined the upstream determinants of ribosome pausing during limitation for arginine and leucine, we sought to investigate its downstream consequences. Towards this goal, we considered the impact of ribosome pausing on cellular translation. We measured global protein synthesis rate during limitation for leucine or arginine by quantifying incorporation of the antibiotic puromycin into nascent polypeptides (Schmidt et al., 2009), and found that global protein synthesis rate was consistently lower during limitation for arginine than leucine (Fig. 5A,B; Supp. Fig. 5A,B). This differential reduction is consistent with similar measurements made previously following extended amino acid limitation (Scott et al., 2000). Based on our previous experiments, we reasoned that three processes could contribute to the regulation of translation during arginine limitation in HEK293T cells: mTORC1 inhibition, GCN2 activation, or ribosome pausing. Given that mTORC1 activity, which stimulates translation initiation, is higher during arginine limitation than leucine limitation (Fig. 3A,C,E; Supp. Fig. 3C,E), mTORC1 signaling cannot account for lower global protein synthesis during arginine relative to leucine limitation.

**Fig. 5.**
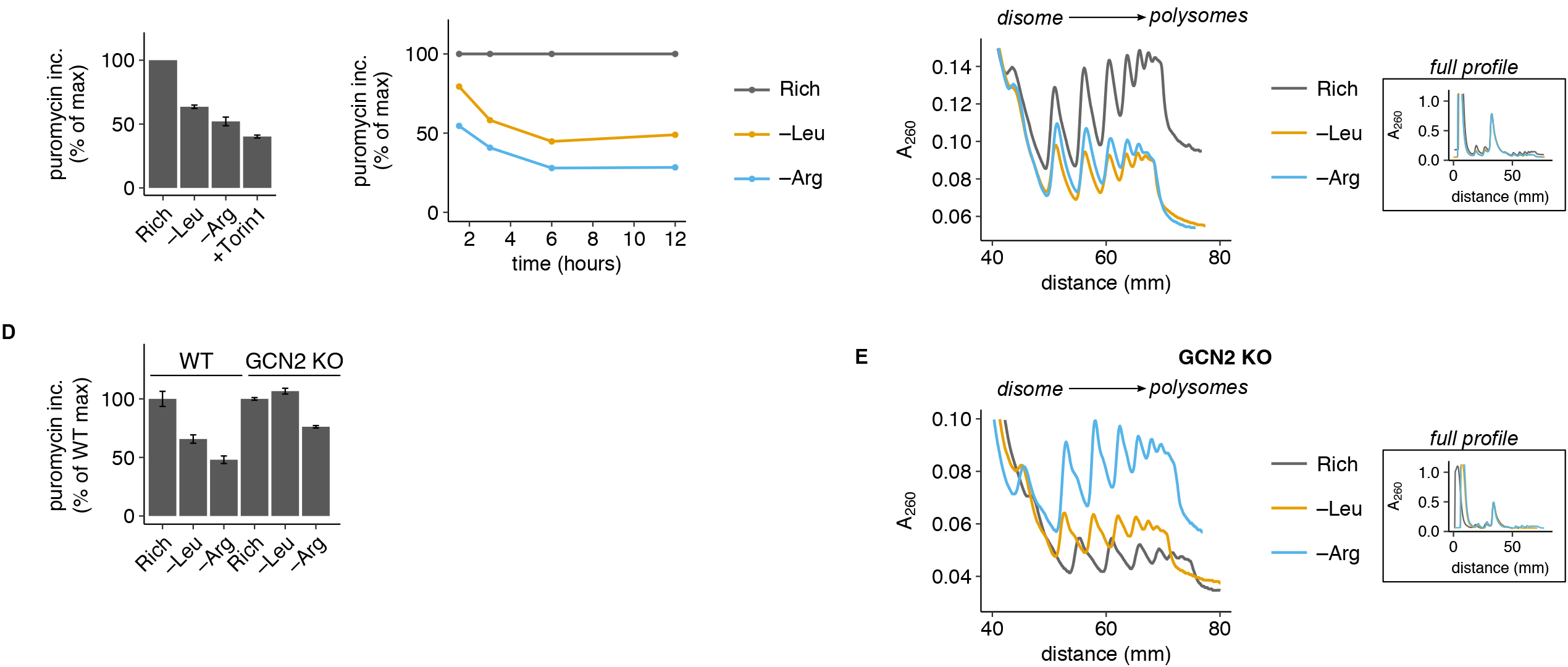
Ribosome pausing reduces global protein synthesis rate during amino acid limitation. **A**) Global protein synthesis rate in HEK293T (WT) cells following 3 hours of leucine or arginine limitation or treatment with 250 nM Torin1, relative to rich medium (calculated as in Supp. Fig. 5A; see Supp. Fig. 5A for representative western blot images). Error bars represent the standard error of the mean for three technical replicate measurements. (**B**) Global protein synthesis rate in HEK293T (WT) cells following 1.5, 3, 6, or 12 hours of leucine or arginine limitation, relative to rich medium (calculated as described in Supp. Fig. 5G). Error bars represent the standard error of the mean for three technical replicate measurements. (**C**) Polysome profiles from WT cells following 6 hours of leucine or arginine limitation or growth in rich medium. The main plot shows overlaid polysome profiles from the disome (2 ribosome) peak to the end of the polysomes for all conditions, the inset plots show the entire profile. All traces were aligned with respect to the monosome peak height along the y-axis and position along the x-axis. (**D**) Global protein synthesis rate in WT or GCN2 KO cell lines following 3 hours of leucine or arginine limitation, relative to rich medium (calculated as in Supp. Fig. 5G, see Supp. Fig. 5G for representative dot blot images). Error bars represent the standard error of the mean for three technical replicate measurements. (**E**) Polysome profiles as described in C in the GCN2 KO cell line following 6 hours of limitation for leucine or arginine or growth in rich medium.

The principal difference between GCN2- and ribosome pausing-mediated control over translation is that GCN2 regulates initiation, while ribosome pausing regulates elongation. To assess whether initiation or elongation rate control accounts for the greater reduction of global protein synthesis rate upon arginine limitation versus leucine limitation, we used polysome profiling to determine the average number of ribosomes per transcript in each condition. If global protein synthesis rate is lower during arginine limitation due to inhibition of translation initiation, there would be fewer ribosomes per transcript upon limitation for arginine compared to leucine. Instead, if global protein synthesis rate is reduced by slow elongation, we would find relatively more ribosomes per transcript upon arginine limitation. While the polysome fraction was reduced by limitation for either leucine or arginine, it was higher during arginine than leucine limitation in HEK293T cells (Fig. 5C,Supp. Fig. 5C), indicating that there are more ribosomes per transcript during arginine versus leucine limitation despite a lower global protein synthesis rate. Thus, elongation rate control, and not initiation rate control, is more likely to account for the greater repression of global protein synthesis rate upon arginine limitation.

mTORC1 inhibition during amino acid limitation reduces global elongation factor activity by phosphorylation of eukaryotic elongation factor 2 kinase (EEF2K) (Leprivier et al., 2013). In theory, this general inhibition of elongation, rather than codon-specific ribosome pausing, could account for a lower, elongation-limited global protein synthesis rate upon limitation for arginine relative to leucine. To assess the role of EEF2K in our measurements of global protein synthesis rate in each condition, we generated an EEF2K knockout cell line (Supp. Fig. 5D,E). Loss of general elongation factor regulation by EEF2K increased global protein synthesis rate upon arginine and leucine limitation by a similar, small margin (Supp. Fig. 5F). Therefore, downregulation of general elongation factor activity cannot account for the greater reduction of protein synthesis upon arginine than leucine limitation, and we instead attribute this difference to elongation rate control by ribosome pausing.

To isolate and quantify the contribution of ribosome pausing to global protein synthesis rate reduction, we made use of the GCN2 KO cell line, which lacks the initiation rate control response to amino acid limitation through both the GCN2 and mTORC1 pathways (Averous et al., 2016; Harding et al., 2000). We reasoned that any residual inhibition of global protein synthesis rate during arginine or leucine limitation in the GCN2 KO cell line would be due to ribosome pausing. Global protein synthesis rate was reduced by 25% during arginine limitation (Fig. 5D, Supp. Fig. 5G). Strikingly, despite this lower global protein synthesis rate, there was a higher polysome fraction during arginine limitation than in rich conditions in this cell line, (Fig. 5E, Supp. Fig. 5C), consistent with our observation of strong ribosome pausing under these conditions (Fig. 4F). Ribosome pausing also develops at a leucine codon in the GCN2 KO cell line (Fig. 4F), and accordingly the polysome fraction was higher upon limitation for leucine than in rich conditions as well (Fig. 5E). However, there was no change to global protein synthesis rate upon limitation for leucine in the GCN2 KO cell line (Fig. 5D), suggesting that global protein synthesis rate reduction in this condition in wild-type cells is primarily mediated by the mTORC1 and/or GCN2 responses. In conclusion, the inverse relationship between global protein synthesis rate and ribosome loading per transcript upon arginine limitation supports a model in which ribosome pausing limits global protein synthesis rate.

### 1.6. Pause-inducing codons in mRNAs reduce protein expression and induce premature termination of translation

Given that ribosome pausing globally reduces protein synthesis, we next investigated whether pausing on mRNAs specifically inhibits production of the encoded protein. Towards this goal, we adapted a protein synthesis reporter in which YFP is fused to an engineered unstable *E. coli* dihydrofolate reductase (DHFR) domain (Han et al., 2014; Iwamoto et al., 2010). In this reporter system, the unstable reporter is rapidly degraded and fluorescence signal only accumulates upon addition of a stabilizing ligand, trimethoprim (TMP). Fluorescence signal upon arginine and leucine limitation correlated well with the global protein synthesis rates we measured in those conditions (Supp. Fig. 6A, left plot, versus Fig. 5B), suggesting that it faithfully reflects the protein synthesis rate of the reporter. To determine the specific effect of ribosome pausing on reporter protein synthesis rate, we constructed a set of codon variant reporters in which either all arginine codons or all leucine codons were swapped to each of the six arginine or leucine codons, respectively (Fig. 6A).

We first determined whether the pause-inducing arginine codons, CGC and CGU (Fig. 1A-C), would reduce reporter protein synthesis rate during arginine limitation. In wild-type HEK293T cells, YFP-DHFR synthesis rate during arginine limitation was reduced to ∼ 60% relative to its value during rich growth when the pause-inducing codons CGC or CGU were used to encode arginine (Fig. 6B, left plot; Supp. Fig. 6B). The CGC codon reduced YFP production specifically during arginine limitation in all three cell lines in which we found ribosome pausing at that codon (Fig. 1A-C), as well as in an alternative reporter construct with different 5’ and 3’ UTR elements (Supp. Fig. 6C). YFP synthesis rate was also reduced by use of the AGA codon (Fig. 6B, left plot; Supp. Fig. 6B), suggesting that pausing might emerge at this codon after the extended duration of arginine limitation that was necessary for detectable accumulation of reporter fluorescence. In the GCN2 KO cell line, in which ribosomes pause at CGC, CGU, and AGA codons (Fig. 4F), use of each of these codons also reduced YFP synthesis rate upon arginine limitation (Figure 6C, left plot). Importantly, there was little difference in the measured protein synthesis rates between the arginine codon variants upon leucine limitation, consistent with the absence of ribosome pausing at arginine codons during leucine limitation (Figure 6B,C; right plots; Supp. Fig. 6B). Similarly, the six leucine codon variants had comparable reductions in YFP synthesis rate upon leucine limitation or arginine limitation (Fig. 6D; Supp. Fig. 6B), consistent with the absence of ribosome pausing at these codons in wild-type cells (Fig. 1A-C). However, YFP synthesis rate was strongly reduced for the UUG codon variant in the GCN2 KO cell line upon limitation for leucine, reflecting the emergence of ribosome pausing at this codon in this condition (Fig. 6E, right plot; Fig. 4F). In all cases, ribosome pausing upon amino acid limitation was sufficient to inhibit reporter protein synthesis.

**Fig. 6.**
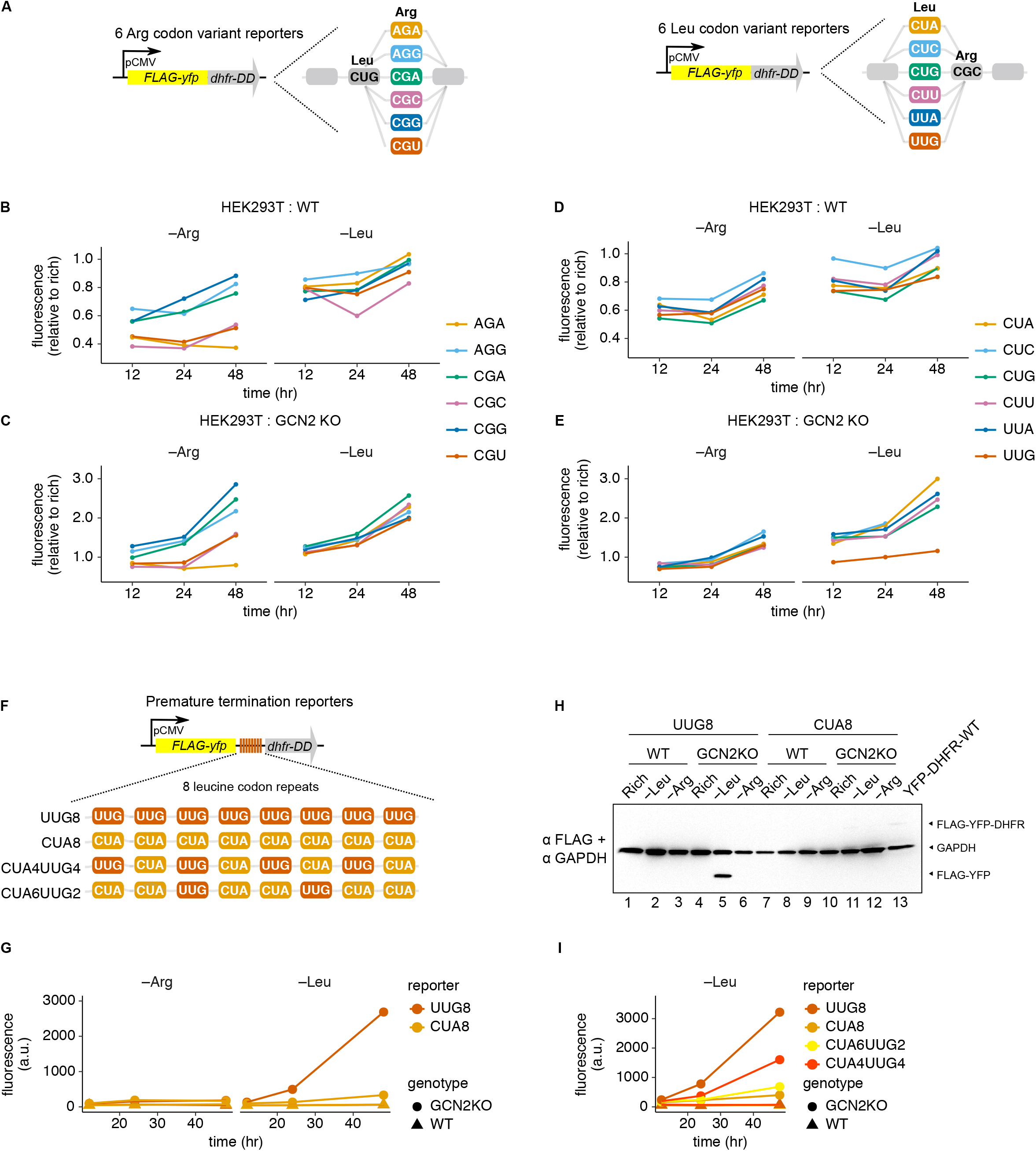
Ribosome pausing reduces protein expression from reporter mRNAs and induces premature termination of translation. (**A**) Arginine and leucine YFP codon variant reporter design. A CMV promoter was used to drive expression of Flag-tagged yellow fluorescent protein (YFP) linked to a dihydrofolate reductase (DHFR) degron domain (DD) (Han et al., 2014), and YFP single codon variants were generated by swapping every arginine or leucine codon to each of the indicated synonymous codons. (**B-E**) YFP fluorescence in the HEK293T (WT) (B,D) or GCN2 KO cell lines (C,E) stably expressing the arginine (B-C) or leucine (D-E) YFP codon variant reporters following limitation for arginine or leucine with 10 µM trimethoprim (+TMP) for 12, 24, or 48 hours, relative to rich medium +TMP. Flow cytometry was used to find the population mean YFP fluorescence for >10,000 events. (**F**) Premature termination reporter design. The reporter described in A was modified by the addition of a short linker of 8 tandem leucine CUA or UUG codons between YFP and DHFR. (**G**) YFP fluorescence in the WT or GCN2 KO cell lines stably expressing the UUG8, CUA4UUG4, CUA6UUG2, CUA8 reporters following limitation for arginine or leucine for 12, 24, or 48 hours without TMP. Flow cytometry was used to find the population mean YFP fluorescence for >10,000 events. (**H**) Western blot probed first for FLAG tag and then for GAPDH after growth in rich medium, or 48 hours of leucine or arginine limitation in the WT or GCN2 KO cell lines stably expressing the UUG8 or CUA8 reporters. Lane 13 contains lysate from the YFP-WT reporter cell line (see Supp. Fig. 4A) for a full-length reporter protein size reference; GAPDH provides an intermediate size reference.

Recent work suggests a role for mRNA degradation in the reduction of protein synthesis rates downstream of slow translation of rare codons in yeast (Presnyak et al., 2015; Radhakrishnan et al., 2016). To determine whether lower reporter protein production rates could be explained by reporter mRNA degradation downstream of ribosome pausing, we measured changes to YFP-CGC and YFP-CGG reporter mRNA levels during arginine and leucine limitation. Levels of YFP-CGC, which contains pause sites, were consistently 2-fold higher than levels of YFP-CGG, which does not contain pause sites, in all conditions (Supp. Fig. 6D). The addition of the reporter stabilizing ligand TMP did not affect mRNA levels (Supp. Fig. 6D). Levels of both YFP-CGC and YFP-CGG were similarly reduced by 50% upon a shift to arginine limitation and unaffected by leucine limitation (Supp. Fig. 6D). Thus, pausing is not clearly linked to a reduction in mRNA levels, and such an effect cannot explain why less protein is produced from the YFP-CGC reporter specifically upon arginine limitation.

To determine whether premature abortive termination might instead account for the reduction of protein synthesis rate by ribosome pausing, as previously described in bacteria (Subramaniam et al., 2014), we modified our protein synthesis rate reporter to detect termination at pause-inducing codons. We inserted a tandem repeat of 8 pause-inducing or non-pause-inducing codons in between the YFP and DHFR domains (Fig. 6F). The full-length YFP-DHFR protein will be degraded efficiently and result in no fluorescence signal. However, abortive termination at the pause-inducing codons would prevent synthesis of the DHFR degron and therefore generate stable YFP. Indeed, we observed a 100-fold increase in YFP fluorescence signal specifically upon leucine limitation when 8 tandem pause-inducing UUG leucine codons (Fig. 4F) were inserted and the reporter (UUG8) was expressed in the GCN2 KO cell line (Fig. 6G). We confirmed that the size of the UUG8 reporter protein corresponded to the predicted size for the premature truncation product in this condition (Fig. 6H). By contrast, we detected only a minor fluorescence increase for the CUA8 reporter upon leucine limitation (Fig. 6G), and the size of the polypeptide produced in this case corresponded to the full length reporter (Fig. 6H). There was no evidence for premature termination of UUG reporter translation in wild-type cells, in which pausing does not occur at UUG codons (Fig. 1A-C), or during limitation for a non-cognate amino acid or in rich conditions. Abortive termination in GCN2 KO cells during leucine limitation correlated positively with the number of pause-inducing codons in the reporter, was detectable when as few as 2 pause sites were present (Fig. 6I), and did not reduce mRNA levels (Supp. Fig. 6E). In fact, abortive termination was associated with increased mRNA level, which may be explained by increased ribosome loading due to stalling upstream of tandem pause sites (Edri and Tuller, 2014). We did not find evidence for similar levels of premature termination at arginine codons during arginine limitation in wild-type cells. This may be because premature termination products are rapidly degraded in these conditions, as polyarginine tracts can trigger ribosome quality control responses (Brandman and Hegde, 2016).

Based on our observation that ribosome pausing reduced protein expression, we sought to identify endogenous proteins whose levels might be regulated by pause-inducing codons during arginine limitation. Towards this goal, we calculated the bias in usage of the pause-inducing arginine codons CGC and CGT for 18,660 coding sequences in the human genome from the genome-wide average usage frequency of these arginine codons (Supp. Fig. 6F).

Among coding sequences biased against use of pause-inducing arginine codons, we found significant enrichment for GO terms broadly related to organelle organization, macromolecule and nitrogen-compound metabolism, RNA processing, and positive regulation of GTPase activity (Supp. Fig. 6G, left plot). Conversely, genes with bias in favor of CGC and CGT codons were significantly enriched for GO terms related to nucleosomes, intermediate filaments, and ion channels involved in neuronal signal transduction (Supp. Fig. 6G, right plot). Given our evidence that ribosome pausing can regulate protein production rates and stimulate premature termination, the genes we identified as being enriched in pause sites are likely to be more translationally repressed upon a shift to arginine-limiting conditions than those depleted of pause sites.

## Discussion

In this work, we investigated how synonymous codons and amino acid availability interact to regulate protein synthesis. We found that ribosome pausing emerges during arginine limitation at two of the six synonymous codons for arginine. We did not find evidence for ribosome pausing at rare codons, or a relationship between pausing and codon optimality or genomic copy number of the cognate tRNA. Instead, it reflected a specific loss of charging for the isoacceptor tRNA(s) that decode those codons. Ribosome pausing developed only in certain environments; ribosomes paused during arginine but not leucine limitation. Rendering these signaling pathways unresponsive to amino acid limitation was sufficient to induce pausing upon leucine limitation, implicating them as upstream determinants of ribosome pausing and thus suggesting that their intrinsic response to arginine limitation is too weak to prevent loss of tRNA charging and the emergence of ribosome pausing. Pausing reduced both global protein synthesis rates as well as expression of specific reporter and endogenous coding sequences. Interestingly, such an effect would not be apparent in ribosome profiling data, as increased ribosome density due to ribosome pausing on a transcript would be associated with reduced protein production. Finally, we found that excessive pausing in the absence of a signaling response to amino acid limitation can result in premature abortive termination at pause-inducing codons.

Despite recent evidence that tRNA level and synonymous codon usage influence translation in mammalian systems (Gingold et al., 2014; Goodarzi et al., 2016; Saikia et al., 2016), we did not find a correlation between ribosome pausing upon arginine limitation and these quantities. Ribosome pausing observed in bacteria is also not explained by these measures (Subramaniam et al., 2013b, 2014). However, an exact accounting of the tRNA supply for each codon is challenging given the degeneracy introduced by wobble decoding, extensive tRNA modifications that influence codon reading, and multiple codons which compete for a single tRNA species. Furthermore, we did not measure tRNA levels but used tRNA gene number to estimate them. Therefore, we cannot exclude the possibility that a more accurate accounting of tRNA supply would explain the observed hierarchy of ribosome pausing. However, it is more likely the balance between tRNA supply and codon usage demand which determines differential isoacceptor sensitivity to changes in arginine levels, as observed in bacteria (Dittmar et al., 2005; Elf et al., 2003). We propose that a consideration of nutrient context is critical for defining which codons or tRNAs are functionally “optimal”.

Our measurements of tRNA charging loss upstream of ribosome pausing suggest that even a 50% charging level for many tRNAs upon amino acid limitation was insufficient to cause ribosome pausing at the cognate codons. This reflects a robustness of ribosome elongation rate to fluctuations in charged tRNA concentrations, and thus changes in charged tRNA concentrations (Saikia et al., 2016) might not always cause changes in translation elongation rate. This finding is also consistent with the proposal that tRNA abundance in mammals is unlikely to be evolutionarily optimized for globally efficient translation (Galtier et al., 2017). Instead, an understanding of what underlies the sensitivity of charging for specific isoacceptor to amino acid levels may reveal the evolutionary forces shaping translation elongation.

Our finding that the mTORC1 and GCN2 pathways respond more potently to limitation for different single amino acids highlights an unusual divergence in their roles, challenging the idea that both pathways act co-ordinately to sense amino acid limitation and appropriately regulate translation rate (Park et al., 2017). The mTORC1 response was clearly non-optimal with respect to preserving arginine homeostasis for protein synthesis: mTORC1 responds more weakly to arginine than leucine limitation, even though arginine becomes more rate-limiting for translation than leucine. Given that direct sensors for arginine (Chantranupong et al., 2016; Wang et al., 2015) and leucine (Wolfson et al., 2016) have been identified in the mTORC1 pathway, this observation is surprising; it suggests that the arginine sensors are unable to optimally sense arginine levels and thereby prevent ribosome pausing at arginine codons, while in contrast the leucine sensor can perform this function. One possibility is that in the context of a tissue or a whole organism, arginine limitation might be typically accompanied by additional cue(s) to stimulate an optimal mTORC1 response, and limitation for only arginine *in vitro* might be insufficient to evoke this response. Investigating the response to arginine limitation *in vivo* will shed light on the role of mTORC1 in regulating arginine consumption.

In contrast to mTORC1, GCN2 — which senses uncharged tRNA — appears to respond optimally; it is equally or more strongly activated during arginine than leucine limitation across all three cell lines. This raises the question of why this robust GCN2 response is insufficient to prevent pausing. It has been recently shown that GCN2 can also sense ribosome pausing, creating a feedback regulation loop between elongation and initiation rates (Ishimura et al., 2016). Therefore, GCN2 activation may in part be downstream of the emergence of ribosome pausing. Dissecting the dynamics of the GCN2 response to amino acid limitation with respect to the emergence of ribosome pausing will clarify whether its role is primarily to prevent, or to respond to, such a loss of amino acid homeostasis. Irrespective of whether it is upstream or downstream of ribosome pausing, the GCN2 response is insufficient to prevent pausing at the timescales explored in this study.

We found that the response through the mTORC1 and GCN2 pathways to single amino acid limitation determines the magnitude of ribosome pausing, presumably by controlling the flux of that amino acid towards anabolic processes and thereby its availability for tRNA charging and translation. This hypothesis is supported by the observation that nonessential amino acids such as glutamine and serine contribute predominately to protein synthesis rather than to cellular metabolite pools in multiple human cell lines (Hosios et al., 2016) As these two signaling pathways regulate multiple facets of cellular metabolism and growth in addition to global translation (Castilho et al., 2014; Laplante and Sabatini, 2013), it is difficult to pinpoint the principal metabolic process that determines the level of each amino acid during its limitation. However, a major role for arginine and leucine flux into translation is supported by our finding that a brief treatment of starved cells with the elongation inhibitor cycloheximide led to recovery of arginine and leucine tRNA charging (Supp. Fig. 4I). Interestingly, inhibition of elongation did not completely rescue tRNA^Arg^_ACG_ charging. As arginine is a nonessential amino acid with several routes for usage in metabolism (Morris, 2007), it is likely that arginine levels during its limitation are also influenced by the flux through these pathways. We expect that perturbing the activity of individual effectors downstream of mTORC1 and GCN2 kinases will identify the contribution of individual metabolic processes to homeostasis of the limited amino acid.

Although we found that the signaling response to amino acid limitation was necessary to prevent ribosome pausing in HEK293T cells, we note that other mechanisms may exert control over ribosome pausing in distinct cell types. For example, high rates of protein catabolism or lysosomal amino acid content could buffer intracellular amino acid levels. Alternatively, a slow cell growth and division cycle or low global protein synthesis capacity could reduce amino acid consumption rates. Indeed, we found no ribosome pausing at leucine codons upon limitation for leucine in HCT116 cells despite a weak amino acid signaling response (Fig. 3C-F, Supp. Fig. 3G). A mechanistic investigation in multiple cell types will clarify the range of cellular processes that exert control over ribosome pausing.

We find that ribosome pausing reduces both global and gene-specific protein synthesis rates. The effects of slow translation at specific codons on protein production have been widely linked to mRNA decay: recent work in yeast has suggested that ribosome stalling at non-optimal codons represses protein synthesis rates by increasing mRNA decay rates (Presnyak et al., 2015; Radhakrishnan et al., 2016). Stalled ribosomes at truncated or damaged RNAs or polybasic sequences are targeted by “no-go decay” (NGD) pathways (Shoemaker and Green, 2012; Simms et al., 2017), which induce endonucleolytic cleavage and degradation of nascent chain polypeptides (Brandman and Hegde, 2016). We did not find evidence for a reduction in mRNA levels due to pausing, although we cannot exclude an increase in mRNA decay rate balanced by an increased synthesis rate. In addition, measurements of total mRNA levels cannot be easily compared to protein production rates because they may include untranslated mRNAs. It is notable, though, that significant changes in mRNA levels have not been observed in cases where protein production is altered by ribosome pausing at specific codons during amino acid limitation (Saikia et al., 2016; Subramaniam et al., 2013b, 2013a, 2014). Perhaps pausing during limitation in the presence of excess uncharged tRNA is qualitatively different from typical “no-go” pauses that result from overall tRNA scarcity, and thus might not stimulate NGD (Buskirk and Green, 2017). We did find evidence for truncated nascent peptides upon ribosome pausing at leucine codons in GCN2 KO cells, suggesting that pausing due to limiting charged tRNA can trigger abortive termination of translation, although the factors involved remain to be elucidated.

Finally, our work raises the question of whether cell-autonomous ribosome pausing is a deleterious, neutral, or an adaptive response. In bacteria, ribosome stalling during amino acid limitation is used as a sensor for upregulating amino acid biosynthesis genes and for entering into a biofilm state, suggesting an adaptive role for ribosome pausing (Dittmar et al., 2005; Subramaniam et al., 2013b). By contrast, in *S. cerevisiae*, an insufficient TOR response to leucine limitation leads to loss of cell viability, and is thus considered to be “non-optimal” (Boer et al., 2008). Analogously, we find that ribosome pausing upon leucine or arginine limitation is linked to a loss of cell viability (Supp. Fig. 7), and it is well known that arginine limitation induces cell death in multiple cancer cell lines (Lind, 2004). Ribosome pausing has also been linked to disease states in mouse and human tissues (Ishimura et al., 2014; Loayza-Puch et al., 2016a). These results suggest that pausing may have a deleterious effect on the cell, for example via protein misfolding or mistranslation stress (Drummond and Wilke, 2008). Pausing might also be a symptom of an upstream loss of metabolic homeostasis; constitutive mTORC1 signaling and elevated consumption of specific amino acids, which according to our findings may synergistically induce ribosome pausing, are characteristic features of certain cancers (Hattori et al., 2017; Jain et al., 2012; Knott et al., 2018; Krall et al., 2016; Loayza-Puch et al., 2016a; Possemato et al., 2011; Scott et al., 2000; Wise and Thompson, 2010). Alternatively, our finding that genes broadly involved in RNA metabolism are biased against the use of arginine pause site codons suggests that ribosome pausing might play a role in metabolic adaptation to arginine limitation, as arginine can contribute to nucleotide synthesis via aspartate (Rabinovich et al., 2015). Further, histone genes are biased towards use of pause sites, and reduced production of nucleosomes could underlie the S-phase cell cycle arrest that accompanies arginine limitation (Nelson et al., 2002; Scott et al., 2000) – though it is unclear whether a prolonged S-phase would be adaptive or detrimental. Therefore, it will be important to determine whether ribosome pausing in this and other contexts plays a positive role in adapting cellular metabolism and gene expression to amino acid limitation, or instead increases cellular stress under these conditions.

## Materials and Methods

### Raw data and code for generation of figures

Full code and detailed instructions for generating the final figures in our paper starting from raw sequencing data is provided as a README.md file and an interactive Jupyter notebook (Perez and Granger, 2007) in the following Github repository (https://github.com/rasilab/adarnell_2018).

### Construction of plasmids

All plasmids and cell lines are included in a key resources table supplementary file.

AAVS1-CAG-hrGFP was from Su-Chun Zhang (Addgene plasmid #52344) (Qian et al., 2014). We cloned sequences for Flag-RagB-WT and Flag-RagB-Q99L into this plasmid in place of hrGFP, from sequences in Flag pLJM1 RagB wt (Addgene plasmid #19313) and Flag pLJM1 RagB 99L (Addgene plasmid #19315) from David Sabatini (Sancak et al., 2008). The resulting CRISPR homology donor plasmids AAVS1-CAG-hrGFP, AAVS1-CAG-RagBWT, and AAVS1-CAG-RagBQ99L were then introduced into HEK293T cells by CRISPR/Cas9 mediated homologous recombination with the AAVS1 sgRNA and Cas9 expression plasmid px330-AAVS1-T2 (see Stable overexpression cell line generation by CRISPR/Cas9 genome editing section). px330-AAVS1-T2 was cloned by inserting the AAVS1-T2 target sequence (GGGGCCACTAGGGACAGGAT) (Mali et al., 2013) into the px330-U6-Chimeric-BB-CBh-hSpCas9 plasmid, from Feng Zhang (Addgene #42230) (Cong et al., 2013) (see Fig. 4C).

To generate sgRNA plasmids for targeting endogenous GCN2 (alias EIF2AK4) and EEF2K (see Supp. Fig. 4G and 5D), sgRNA sequences were obtained from a list of validated guides from the 3^rd^ generation lentiGuide-Puro library (Doench et al., 2016). Two sgRNA sequences each targeting exonic sequences ∼790 bp apart in GCN2 (from Addgene plasmids #75876 and 75877), and ∼230 bp apart in EEF2K (from Addgene plasmids #77855 and 77856), were selected. For each pair targeting a gene, one sgRNA was cloned into pU6-(BbsI)_CBh-Cas9-T2A-BFP, from Ralf Kuehn (Addgene plasmid #64323) (Chu et al., 2015), and the other was cloned into pSpCas9(BB)-2A-GFP (PX458), from Feng Zhang (Addgene plasmid #48138) (Ran et al., 2013) using T4 DNA ligase (NEB) after BbsI digestion. This produced the targeting sgRNA-Cas9 plasmids pU6-GCN2-1-Cas9-2A-BFP, pU6-GCN2-2-Cas9-2A-GFP, pU6-EEF2K-1-Cas9-2A-BFP, and pU6-EEF2K-2-Cas9-2A-GFP. For generation of knockout cell lines, see Knockout cell line generation by CRISPR/Cas9 genome editing section.

Our YFP-DHFR protein synthesis rate reporters (see Fig. 6A) were built by cloning from pLJM1-EGFP, from David Sabatini (Addgene plasmid #19319) (Sancak et al., 2008). The EGFP coding sequence in this vector was replaced by the YFP-DHFR sequence from KHT61-Unreg-YFP-DD, a gift from Kyuho Han (Han et al., 2014)), along with an N-terminal Flag epitope tag to generate the base “wild-type” (YFP-WT) pLJM1-Flag-YFP-DHFR reporter was cloned into pLJM1 in place of EGFP. The YFP-WT reporter has 13 CGC and 1 CGU arginine codons, and 23 CUG, 5 CUC, 2 UUA, and 2 UUG leucine codons. To generate codon variants, synthetic DNA gBlocks (IDT) were ordered in which all 14 arginine codons in YFP and DHFR, or all 21 leucine codons in YFP were swapped to one out of each of the six synonymous arginine or leucine codons; these gBlocks were amplified by PCR and cloned in place of the YFP-WT sequence in the pLJM1 plasmid backbone. The following library of Flag-tagged codon variant reporter lentiviral donor plasmids was generated: YFP-WT, YFP-CGG, YFP-CGA, YFP-CGU, YFP-AGA, YFP-AGG, YFP-CUA, YFP-CUC, YFP-CUU, YFP-UUA, and YFP-UUG. These plasmids were used to generate stable reporter cell lines by lentiviral transduction into HEK293T, HeLa, HCT116, and the HEK293T GCN2 KO cell line (see Stable overexpression cell line generation by lentiviral transduction section).

The YFP-DHFR protein synthesis rate reporters were modified to generate premature termination reporters (see Fig. 6F) by cloning in eight tandem leucine codons into the pLJM1-YFP-CUA lentiviral donor plasmid in between the YFP and DHFR sequences. YFP and DHFR were amplified by PCR with leucine codons added in the reverse primer overhang sequences for YFP and forward primer overhang sequences for DHFR, and these sequences were then re-assembled into the pLJM1 backbone. The following library of four Flag-tagged premature termination reporter lentiviral donor plasmids was generated, in which the codon and following number refer to the composition of the eight tandem leucine codon repeat: UUG8, CUA8, CUA4UUG4, CUA6UUG2. These plasmids were used to generate stable reporter cell lines by lentiviral transduction into HEK293T and the HEK293T GCN2 KO cell line (see Stable overexpression cell line generation by lentiviral transduction section).

A variant YFP-DHFR protein synthesis rate reporter (see Supp. Fig. 5B) was built by cloning from pAAVS1P-iCAG.copGFP, from Jizhong Zou (Addgene plasmid #66577) (Cerbini et al., 2015). To generate pAAVS1P-iCAG.FlagYFP-DHFR-WT and –CGG codon variant reporters, sequences from pLJM1-Flag-YFP-DHFR-WT (YFP-WT) and YFP-CGG were cloned in place of copGFP. These plasmids were used as homology donors to generate stable reporter cell lines in HEK293T cells by CRISPR/Cas9-mediated homologous recombination with the AAVS1 sgRNA and Cas9 expression plasmid px330-AAVS1-T2 (as for the AAVS1-CAG-hrGFP and its derivative plasmids described above; see Stable overexpression cell line generation by CRISPR/Cas9 genome editing section).

### Cell culture and amino acid limitation

HEK293T, HeLa, and HCT116 adherent cells (HEK293T and HeLa obtained from ATCC, catalog numbers CRL-3216 and CCL-2; HCT116 obtained from the National Cancer Institute (NCI) panel of 60 cancer cell lines) were passaged in high-glucose DMEM without pyruvate (Gibco) with penicillin/streptomycin (Corning) and 10% fetal bovine serum (FBS; ATCC catalog number 30–2020). Amino acid limitation media were prepared from low glucose DMEM powder without amino acids (US Biological catalog number D9800–13) according to manufacturer’s instructions; all amino acids except leucine and arginine, and glucose were supplemented according to this recipe: 3 g/L additional glucose, 30 mg/L glycine, 63 mg/L cysteine 2·HCl, 580 mg/L glutamine, 42 mg/L histidine HCl·H_2_O, 105 mg/L isoleucine, 146 mg/L lysine HCl, 30 mg/L methionine, 66 mg/L phenylalanine, 42 mg/L serine, 95 mg/L threonine, 16 mg/L tryptophan, 64 mg/L tyrosine 2·Na 2·H_2_O, and 94 mg/L valine. Media was subject to vacuum filtration and then supplemented with 10% dialyzed FBS (Invitrogen catalog number 26400-044) before use. For all amino acid limitation assays – except time course experiments over multiple days (see sections for Flow cytometry and Cell viability) – cells were expanded to 75% confluency, washed once in PBS, and transferred to limitation medium supplemented with either leucine (for arginine limitation) or arginine (for leucine limitation), or both (for rich medium).

For all experiments, technical replicates refer to the repetition an entire experiment with a separate dish of cells split off from the same parental cell line (i.e. produced from the same lentiviral transduction or CRISPR editing process).

### Stable overexpression cell line generation by lentiviral transduction

HEK293T cells were transfected at 75% confluency in a 10 cm plate with donor expression plasmid pLJM1 containing the desired insert, and the lentiviral packaging plasmids psPAX2, from Didier Trono (Addgene plasmid #12260), and pCMV-VSV-G, from Bob Weinberg (Addgene plasmid #8454) (Stewart et al., 2003) in a 10:9:1 ratio (by weight) using Lipofectamine 3000 (Invitrogen), according to the manufacturer’s instructions. The media was replaced after 12–16 hours, and lentivirus was harvested at 48 hours by passing culture supernatant through a low-protein binding filter with 0.45 µm pore size. 1 mL of virus was used to transduce 50–60% confluent cells in a 6 cm plate. Cells were passaged to a 10 cm plate after 24 hours, and antibiotic selection was performed after 48 hours by adding puromycin (2 µg/ml for HEK293T cells, 1 µg/ml for HCT116, HeLa cells). Cells were passaged in selection media for 2–4 days, until non-transduced cells treated with puromycin in a parallel plate were fully dead, and were then expanded for generation of stocks and experiments.

### Stable overexpression cell line generation by CRISPR/Cas9 genome editing

All transfections were performed at 75% confluency using Lipofectamine 3000, according to manufacturer’s instructions.

To generate hrGFP, RagB-WT, and RagB-Q99L cell lines (see Fig. 4C): HEK293T cells in a 6 well plate were transfected with homology donor plasmid (pAAVS1-CAG-hrGFP, pAAVS1-CAG-RagBWT, or pAAVS1-CAG-RagBQ99L) and the px330-AAVS1-T2 guide RNA plasmid at a ratio of 4:1 (2 µg donor : 500 ng guide). Homologous recombination and expression of the hrGFP fluorescent protein or FLAG-tagged RagB transgenes was confirmed in the resulting polyclonal population by PCR, flow cytometry, and western blotting after puromycin selection (as described in “Stable overexpression cell line generation by lentiviral transduction”).

To generate arginine/leucine codon variant YFP-DHFR reporter cells lines (see Supp. Fig. 6B): HEK293T cells in a 10 cm plate were transfected with homology donor plasmid (for YFP reporter lines: pAAVS1P-iCAG.copGFP, pAAVS1P-iCAG.Flag-YFP-DHFR-WT, pAAVS1P-iCAG.Flag-YFP-DHFR-CGG) and px330-AAVS1-T2 guide RNA plasmid at a ratio of 2:1 (10 µg donor : 5 µg guide). Homologous recombination and TMP-inducible YFP fluorescence were confirmed in the resulting polyclonal population by PCR, flow cytometry, and western blotting after puromycin selection (as described in “Stable overexpression cell line generation by lentiviral transduction”).

### Knockout cell line generation by CRISPR/Cas9 genome editing

75% confluent HEK293T cells in a 12 well plate were transfected with 500 ng of each targeting RNA plasmid, using Lipofectamine 3000 according to the manufacturer’s instructions, in the following four combinations: 1) both pU6-GCN2-1-Cas9-2A-BFP and pU6-GCN2-2-Cas9-2A-GFP, 2) both pU6-EEF2K-1-Cas9-2A-BFP and pU6-EEF2K-2-Cas9-2A-GFP, 3) pU6-GCN2-1-Cas9-2A-BFP only, and 4) pU6-GCN2-2-Cas9-2A-GFP only. Cells were transferred to a 6 well plate 24 hours post transfection. Single fluorescent (BFP+/GFP+) cells were sorted into individual wells of a 96 well plate by FACS. 96 well plates with isolated clones were spun 100xG for 1 minute to sediment cells. Clones were allowed to expand for 14 days and then passaged for generation of stocks and Western blot analysis to confirm complete knockout of GCN2 or EEF2K. 92% of clones tested in this manner were positive for complete GCN2 KO (11/12), 83% were positive for EEF2K KO (10/12) (see Supp. Figs 4G and 5D).

### Ribosome profiling

To detect codon-specific ribosome pausing, ribosome profiling was performed according to the following protocol (Ingolia et al., 2009), with modifications detailed below (see Fig. 1,4 and Supp. Fig. 1,4).

Cells were expanded to 75% confluency in two 15 cm plates harvested for ribosome profiling. Cells were washed once, briefly, in ice cold PBS. PBS was thoroughly drained, and plates were immediately immersed in liquid nitrogen for flash freezing and then transferred to –80C. Frozen cells were lysed on each plate by scraping into 300 µL lysis buffer (20 mM Tris pH 7.5, 15 mM MgCl_2_, 150 mM NaCl, 100 µg/mL cycloheximide, 5 mM CaCl_2_, 1% Triton, 50 U/mL Turbo DNase), and lysates from the two 15 cm plates were combined to yield ∼1 mL of lysate. Ribosome footprints were generated from 450 µL of lysate by 1 hour of digestion with 800 U micrococcal nuclease (MNase, Worthington Biochemical) at room temperature (25˚C) with nutation, which was quenched by addition of 4.5 µL 0.5 M EGTA. Footprints were purified by sucrose density gradient fractionation; a BioComp Gradient Station was used to generate 10–50% sucrose density gradients (Seton Polyclear 14×89 mm tubes) in 1X polysome resuspension buffer (20 mM Tris pH 7.5, 15 mM MgCl_2_, 150 mM NaCl, 100 µg/mL cycloheximide). 400 µL MNase digested lysate were loaded onto gradients in SW41 rotor buckets (Beckman Coulter) after removing 260 µL of the gradient from the top, and samples were ultracentrifuged in an SW41 rotor for 2.5 hours at 35,000 RPM and 4˚C (Beckman Coulter). Fractionation was performed at 0.22 mm/sec with UV absorbance monitoring at 254 nm (EconoUV Monitor) and the monosome fraction was collected in addition to the contiguous disome “shoulder” (∼2.5 mL in total). Total RNA was purified from sucrose solution by addition of 7 mM EDTA and 1% SDS, extraction in Acid-Phenol:Chloroform pH 4.5 with isoamyl alcohol at 25:24:1 (Invitrogen) at 65˚C, and extraction in chloroform. RNA was precipitated by addition of 1/9^th^ volume 3M NaOAc pH 5.5, 2 µL Glycoblue (Applied Biosystems), and isopropanol to the aqueous supernatant.

Ribosome footprints were purified by loading 8 µg of the gradient fraction RNA on 15% TBE-Urea gel (Bio-Rad) and electrophoresed at 200V for 65 minutes alongside the 3’ phosphorylated 26 nt RNA NI-NI-20 (Ingolia et al., 2012) and low range ssRNA ladder (NEB) as size standards, gel was stained in SYBR Gold, and footprints were excised in a wide range from ∼26–40 nt. Gel slices were passed through a 0.6 mL tube with a needle hole in the bottom nested in a 1.5 mL tube to create a gel slurry, and RNA was extracted in 0.3 M NaOAc pH 5.5, 1 mM EDTA, and 100 U/mL Superase-In (Invitrogen) overnight at room temperature with rotating and then precipitated by addition of 2ul glycoblue and isopropanol.

Footprints were dephosphorylated with T4 PNK (NEB) according to the manufacturer’s instructions for 1 hour at 37˚C, then precipitated. Footprints were then polyA-tailed with E. coli polyA polymerase (NEB) according to the manufacturer’s instructions for 10 minutes at 37˚C, then precipitated. Reverse transcription was performed using SuperScriptIII (Invitrogen) and 0.5 µM oNTI19pA oligo primer (Ingolia et al., 2009) for 30 minutes at 48˚C, and RT products were purified by running a 10% TBE-Urea gel at 200V for 65 minutes, using a “no template” sample as a size standard for the RT primer alone. RT products were purified from gel slices using the approach described above for ribosome footprints (in 0.3 M NaCl, 1 mM EDTA, and 0.25% SDS) and then precipitated. RT products were circularized with CircLigase (Epicentre) for 60 minutes at 60˚C, then precipitated. rRNA was removed by subtractive hybridization with MyOne Streptavidin Dynabeads; biotinylated reverse complement oligos to two discrete rRNA sequences that were recovered extremely abundantly in our test ribosome profiling libraries (o3285, o3287) were annealed to circularized libraries in a Thermocycler, beads were prepared according to (Ingolia et al., 2012) and an equal volume was added to annealed oligo/libraries for 15 minutes at 37˚C. Supernatant was recovered and precipitated. Resulting libraries were amplified by 6–12 cycles of PCR with common (reverse) and unique 6nt index (forward) library primers and purified after running on a 10% TBE gel at 200V for 60 minutes. ∼170 nt dsDNA libraries were extracted from gel slices using same method as for RT products, precipitated, resuspended in 10 µL Tris 10 mM pH 7, and quantified using a TapeStation or BioAnalyzer instrument. Up to 15 multiplexed libraries were submitted for sequencing on both lanes of an Illumina HiSeq 2500 Rapid Flow Cell at 3–4 nM in <10 µL. Sequencing runs yielded approximately 150 million reads in total for all multiplexed libraries.

Notably, two ribonucleases, RNase I and micrococcal nuclease (MNase), are commonly used for ribosome profiling (MGlincy and Ingolia, 2017). We analyzed monosome-bound RNA footprint generation by these enzymes using sucrose density gradient fractionation. We observed near-complete degradation of the 60S ribosomal subunit and ribosome-bound mRNA fractions by RNase I in buffers with either high (Ingolia et al., 2012) or low magnesium (Andreev et al., 2015) and across a broad range of RNase I concentrations (Supp. Fig. 1A,B), similar to results obtained in *Drosophila* (Dunn et al., 2013). In contrast, the 60S and monosome fractions were largely intact after digestion with MNase (Supp. Fig. 1C), and therefore we used this nuclease to generate monosome-bound RNA footprints, which were then purified by sucrose density gradient fractionation and size selection (Supp. Fig. 1D) before sequencing. As previously reported (Dunn et al., 2013; Reid et al., 2015), MNase results in slightly longer reads and a broader read length distribution (Supp. Fig. 1E) as it does not digest completely around bound ribosomes. However, read density exhibited robust three nucleotide periodicity (Supp. Fig. 1G, lower panel), is clearly enriched in the coding region, and exhibits peaks at start and stop codons (Supp. Fig. 1F,G), allowing resolution of codon-level changes in translation elongation.

### Ribosome profiling data analysis

Analysis was performed using R and Bash programming languages. Full code and detailed instructions for generating the final figures in our paper starting from raw sequencing data is provided as a README.md file and an interactive Jupyter notebook (Perez and Granger, 2007) in the following Github repository (https://github.com/rasilab/adarnell_2018).

The polyA tail was trimmed from 50 nt single-end raw sequencing reads using `cutadapt`(Martin, 2011) with a minimum length cutoff of 13 nt. A subtractive alignment was performed against ribosomal RNA using `bowtie` (Langmead et al., 2009), and the remaining reads were aligned to a transcriptome index using `rsem` and `bowtie` (Li and Dewey, 2011). To calculate the pre-processing statistics and assess library quality (Supp. Fig. 1E-G), we used 3’ trimming of 12 nt for reads <= 32 nt and 13 nt trimming for reads > 32 nt to demonstrate 3 nt periodicity. However for the rest of the analyses, since we were interested in the overall increase in ribosome occupancy at codons and frame information was not required for this analysis, we trimmed 12 nt from both sides to smooth our ribosome density profiles as described in previous MNase-based studies in bacteria (Li et al., 2012; Oh et al., 2011; Subramaniam et al., 2014). To calculate reads counts for each transcript, each transcript position aligning to the trimmed read was assigned a count of the inverse value of the read length. The DESeq2 package was used to normalize each sample and then calculate gene fold changes (see for example Fig. 3C,D) (Love et al., 2014).

To calculate the average ribosome occupancy around each codon, only transcripts with a minimum read density of 1 read per codon were considered. Reads at each transcript position were first normalized to the mean read count for that transcript. For each codon, the average read coverage was found for each position in a 150 nt window on either side of all occurrences of that codon.

To calculate the change in average ribosome occupancy around each codon upon amino acid limitation (see for example Fig. 1B), the average ribosome occupancy at each position in the 150 nt window around the codon in rich conditions was subtracted from that in an amino acid limited condition. To calculate the summed ribosome occupancy at each codon (see for example Fig. 1A), this 300 nt average ribosome occupancy vector for each codon was summed.

### Polysome profiling

The same procedure as in the “Ribosome profiling” section was used, with the following modifications (see Fig. and Supp. Fig. 5). Nuclease digestion was excluded, and 150 µL of clarified lysate was loaded directly onto sucrose density gradients. Gradients were centrifuged in a SW41 rotor at 35,000 RPM for 3 hours at 4˚C with the “slow” brake setting (Beckman Coulter). Polysome profiles were analyzed by fractionation at 0.22 mm/second using the BioComp Gradient Station and Gradient Profiler software, with UV monitoring at A254 nm (EconoUV). The relative polysome to monosome fraction area was calculated for each profile by 1) subjective definition of the fraction boundaries, 2) subtracting the lowest value in the profile from all points along the profile, and 3) manual integration using the trapezoid rule (see Supp. Fig. 5C).

### tRNA charging analysis

tRNA charging analysis was performed according to (Varshney et al., 1991) with the following modifications (see Fig. 2 and Supp. Fig. 4). 75% confluent cells in a 10 cm plate were washed once in PBS and flash frozen. Cells were scraped into ice cold 500 µL AE buffer (0.3 M NaOAc pH 4.5, 10 mM EDTA) on plates and added to 500 µL ice cold acid-saturated phenol:chloroform pH 4.5 (with isoamyl alcohol, 125:24:1, Invitrogen). Extractions were vortexed hard for 10 minutes, rested on ice for 3 minutes, and spun for 10 minutes at 20,000xG at 4˚C. Aqueous supernatant was recovered and precipitated by adding 2 µL glycoblue and isopropanol. The pellet was resuspended in 10 mM NaOAc pH 4.5, 1 mM EDTA. RNA was deacylated in 100 mM Tris pH 9 at 37˚C for 30 minutes, then precipitated and resuspended in 10 mM NaOAc pH 4.5, 1 mM EDTA as a control for electrophoretic mobility of uncharged tRNA.

For acid urea gel electrophoresis, 500 ng – 1 µg RNA and deacylated control in 0.1 M NaOAc pH 4.5, 8 M urea, 0.05% bromphenol blue, and 0.05% xylene cyanol were electrophoresed on a 0.4 mm 6.5% polyacrylamide gel with 8M urea in 0.1M NaOAc pH 4.5 at 450V and 4˚C for 18–20 hours. The gel region between the loading dye bands was excised and transferred according to “Northern blotting” section.

Probes were designed to hybridize uniquely to tRNA isoacceptors, where possible, or isodecoders after alignment of all arginine and leucine tRNAs (sequences from the Genomic tRNA database http://gtrnadb.ucsc.edu/ (Chan and Lowe, 2016); alignment performed using Muscle (Edgar, 2004)). tRNAs with introns and psueo-tRNAs were identified using the tRNAscan-SE program (http://lowelab.ucsc.edu/tRNAscan-SE/) (Lowe and Eddy, 1997). All probes were validated for specificity by Northern blotting against in vitro transcribed target tRNAs and equimolar amounts of the most likely tRNA candidate for cross-hybridization. Candidate cross-hybridizing tRNAs were identified by a genomic tRNA BLAST. We were not able to find uniquely hybridizing probe for tRNA^Leu^_AAG_ and tRNA^Leu^_UAG_ as these leucine isoacceptor genes have a great degree of sequence homology; however the major species detected for the AAG and UAG probes is the indicated tRNA.

### Western blotting

75% confluent cells in a 10 cm plate were lysed by scraping and pooling in 300 µL of 50 mM HEPES pH 7.4, 40 mM NaCl, 2 mM EDTA, 1 mM sodium orthovandate, 10 mM sodium glycerophosphate, 10 mM sodium pyrophosphate, 50 mM sodium fluoride, 1% Triton X-100. After 10 minutes at 4°C, the insoluble fraction was cleared by centrifugation for 10 minutes at 4˚C and 20,000g. Lysate was electrophoresed in 1X SDS sample buffer (BioRad) on a 4–20% Tris Glycine gel (Novex) and blotted onto 0.45 µm nitrocellulose. Primary antibodies (Cell Signaling Technology, CST) from rabbit against GCN2 (3302S), eEF2K (3692S), eEF2 (2332S), P∼T56 eEF2 (2331S), eIF2α (5324P), P∼S51 eFI2α (3398P), S6K (9202S), P∼T389 S6K (9205S), RPS6 (2217S), P∼S235/236 RPS6 (4858S), GAPDH (2118S). Primary antibody from mouse against FLAG (Sigma-Aldrich, F3165) were used at 1:1000. The primary antibody from rabbit against puromycin was used at 1:25,000 (Sigma-Aldrich, MABE343). HRP-conjugated secondary antibodies were used at 1:5000 (anti-rabbit from CST, 7074S; anti-mouse from Sigma-Aldrich, 12-349). 5% BSA (CST) in TBST was used for all blocking and antibody solutions for phospho-antibody blots, and 5% milk in TBST was used for all others. SuperSignal West Femto Substrate (ThermoFisher) was used for developing, and Restore Western Blot stripping buffer (Thermo Scientific) was used to strip blots (see Fig. and Supp. Fig. 3–6).

For dot-blotting, 2 µL of lysate was spotted onto a 0.45 µm nitrocellulose membrane, allowed to dry for 15 minutes, and then blots were processed as described above for western blotting (see Fig. and Supp. Fig. 5).

### Northern blotting

RNA samples were run on 10% TBE-Urea gels (Criterion) or homemade acid-urea polyacrylamide gels (for tRNA charging analysis). Gels were rinsed thoroughly in 0.5X TBE and transferred to HyBond Nylon+ membrane in 0.5X TBE using a semi-dry transfer apparatus at 3 mA/cm^2^ for 1 hour. The blot was crosslinked using the Stratalinker “auto-crosslink” setting once on each side, prehybridized in PerfectHyb (Sigma) buffer for 1 hour at 64˚C, and hybridized at 64˚C with 5 pmol probe. Probes were end-labelled with T4 PNK using [γ-P^32^]-ATP and purified with G25 sepharose columns (GE Healthcare Life Sciences). The blot was washed 2x in a low-stringency wash buffer (2X SSC, 0.1% SDS) and 1X in a high stringency wash buffer (0.5X SSC, 0.1% SDS) at 64˚C, exposed to a Phosphor-Imaging screen for 12 – 24 hours, and imaged using a Typhoon scanner (see Fig. and Supp. Fig. 2,4).

### Flow cytometry

These assays were performed over 12, 24, or 48 hours post-limitation for arginine, leucine, or growth in rich conditions; therefore cells grew to varying degrees of confluency, and initial seeding number was adjusted so that cells grown in nutrient rich conditions would be ∼75% confluent at the time of collection for flow cytometry measurements. Cells in amino acid limited conditions were less confluent. Cells (HEK293T, HCT116, or HeLa) were detached from a 6 or 12 well plate using 0.05% trypsin + EDTA (Invitrogen). Trypsinization was quenched with DMEM + 10% FBS, and cells were pelleted by centrifugation at 125g for 5 minutes. Pellets were resuspended in 500 µL (for a 12 well plate well) to 1 mL (for a 6 well plate well) of PBS and the cell suspension was passed through a 0.35 µm nylon mesh strainer-top tube (Corning) and kept at room temperature for flow cytometry analysis within 1 hour of filtration. 10,000–30,000 events were collected for all experiments. YFP fluorescence measurements were log-transformed and the mean and standard deviation of all events was calculated from the population (see Fig. and Supp. Fig. 6).

### Puromycin incorporation assays

75% confluent cells in a 6 cm plate were limited for arginine or leucine or grown in nutrient rich conditions for the desired time, followed by addition of puromycin (Sigma-Aldrich, P8833) to the culture medium at 10 µg/mL for 5 minutes at 37˚C. After exactly 5 minutes, cells were washed once in ice-cold PBS and flash frozen in liquid nitrogen. Western blots or dot blots were performed to quantify puromycin incorporation into nascent polypeptide chains (see Fig. and Supp. Fig. 5).

### S-35 pulse assay

75% confluent cells in a 6 well plate were limited for arginine or leucine or grown in nutrient rich conditions for the desired time, and then 50 uCi EasyTag ^35^S-labeled methionine (Perkin Elmer) was added to cultures for 30 minutes at 37˚C. Cells were lysed and collected as in “Western blotting” section. 25 µL lysate was spotted onto cellulose acetate filters (Whatman) and dried for 15 minutes. Filters were washed in a glass dish: 1X for 5 minutes in cold 5% TCA, 2X for 5 minutes in cold 10% TCA, 2X for 2 minutes in cold EtOH, and 1X for 2 minutes in cold acetone. Filters were then air dried for 15 minutes and transferred into a scintillation vial with 5 mLs scintillation fluid (ReadySolv-HP) for counting (see Supp. Fig. 5B). Notably, we could not deplete intracellular methionine pools by limitation for methionine, as this would significantly interfere with the amino acid limitation response measured in our experiments.

Problematically, we found that ^35^S-methionine incorporation was higher after 3 hours of limitation for leucine than growth in rich conditions (Supp. Fig. 5B). This is likely an experimental artifact since the uptake rate and intracellular pool size of radiolabelled methionine can change significantly in response to amino acid limitation. We therefore used puromycin incorporation to quantitatively compare protein synthesis rates in subsequent experiments.

### Reverse transcription & qPCR

Reverse transcription using a dT-20 primer (Invitrogen) or gene-specific primers was performed using Superscript III (Invitrogen) according to the manufacturer’s instructions. cDNA template was diluted in water and qPCR was performed in 10 µL reaction volumes in 96 well plates, using the PowerUp SYBR Green PCR master mix according to the manufacturer’s instructions (Thermo Fisher Scientific) (see Supp. Fig. 6C,D). To calculate relative YFP reporter mRNA levels, the YFP C_t_ value from qPCR analysis in each condition was normalized to the GAPDH C_t_ value to find ΔC_t_, and then to the ΔC_t_ for the arbitrary normalization sample (for Fig. 6C, YFP-WT in rich medium; for Fig. 6D, WT-CUA8 in rich medium) to find ΔΔC_t_, which was converted to a normalized mRNA level by taking 2^−ΔΔCt^.

### Cell viability assays

20,000 cells were seeded in 96 well plates (1 plate per assay time point, 5 technical replicates per plate) in amino acid limitation medium or rich medium. At desired time points, CellTiterGlo (CTG) assay (Promega) was performed according to the manufacturer’s instructions with the following modifications. Cells were lysed by adding 1 volume of CTG reagent and then transferred to an opaque black 96 well plate (Perkin Elmer catalog number 6005660) for luminescence reading. Luminesence was measured immediately on a TopCount instrument (Perkin Elmer) at 30°C. All viability measurements were normalized to an initial reading for each well taken 1.5 hours after seeding adherent cells (see Supp. Fig. 7).

### Databases utilized

A subset of unique canonical transcripts used for mapping aligned ribosome profiling sequencing reads was defined based on the Gencode v24 database annotation file (gencode.v24.annotation.gff3). For each gene, only transcripts annotated as both CCDS in the APPRIS principal splice isoform database (Rodriguez et al., 2013) were included; of this subset, the transcript with the lowest CCDS number for each gene was selected to generate a unique set.

tRNA gene numbers (see Supp. Fig. 1K) were obtained from the genomic tRNA database (gtRNAdb; gtrnadb.uscs.edu/Hsapi19/) (Chan and Lowe, 2016).

### Estimation of usage bias for pause-inducing arginine codons and GO analysis

We employed a binomial probability distribution to estimate the probability, for each gene, of having the observed number of CGC and CGU codons given the genome-wide average arginine codon usage frequencies (see Supp. Fig. 1J). To avoid skew due to local GC bias in our analysis, we only considered sets of pause-inducing or non-pause-inducing arginine codons with equivalent GC content (CGC/CGU vs. CGA/CGG, respectively; “CGN codons”). We calculated the average expected number of pause-inducing codons for each gene as the mean of a theoretical binomial probability distribution (µ); n*p, where n is the total number of arginine codons and p is the average frequency of stall sites relative to other CGN codons (∼ 0.46). We also calculated the standard deviation of that theoretical binomial probability distribution (σ) for each gene as the square root of n*p*(1-p). To then calculate a Z-score, we subtracted µ from the observed number of pause-inducing codons in that gene, and normalized by σ. When ranked, the resulting Z-scores represent bias towards (high Z-scores) or against (low Z-scores) the use of pause-inducing arginine codons to encode arginine in each gene (see Supp. Fig. 6E).

Gene ontology (GO) analysis to detect enrichment for GO terms in genes with biased usage of pause-inducing arginine codons was performed in R using the topGO library (Alexa and Rahnenfuhrer, 2016; Grossmann et al., 2007). Full code for generating the final figures in our paper starting from a ranked list of Z-scores (see Estimation of usage bias for pause-inducing codons section) is provided both as an interactive Jupyter notebook and as a static HTML file (Data S3). Fisher’s exact test was used to determine significance in enrichment of GO terms in genes with the highest and lowest 5% of Z-scores. GO terms with a false-discovery rate adjusted p-value of < 0.05 were visualized using R scripts to plot generated by REVIGO (Supek et al., 2011) (see Supp. Fig. 6F).

## Supplementary Figure Legends

**Supp. Fig. 1.**
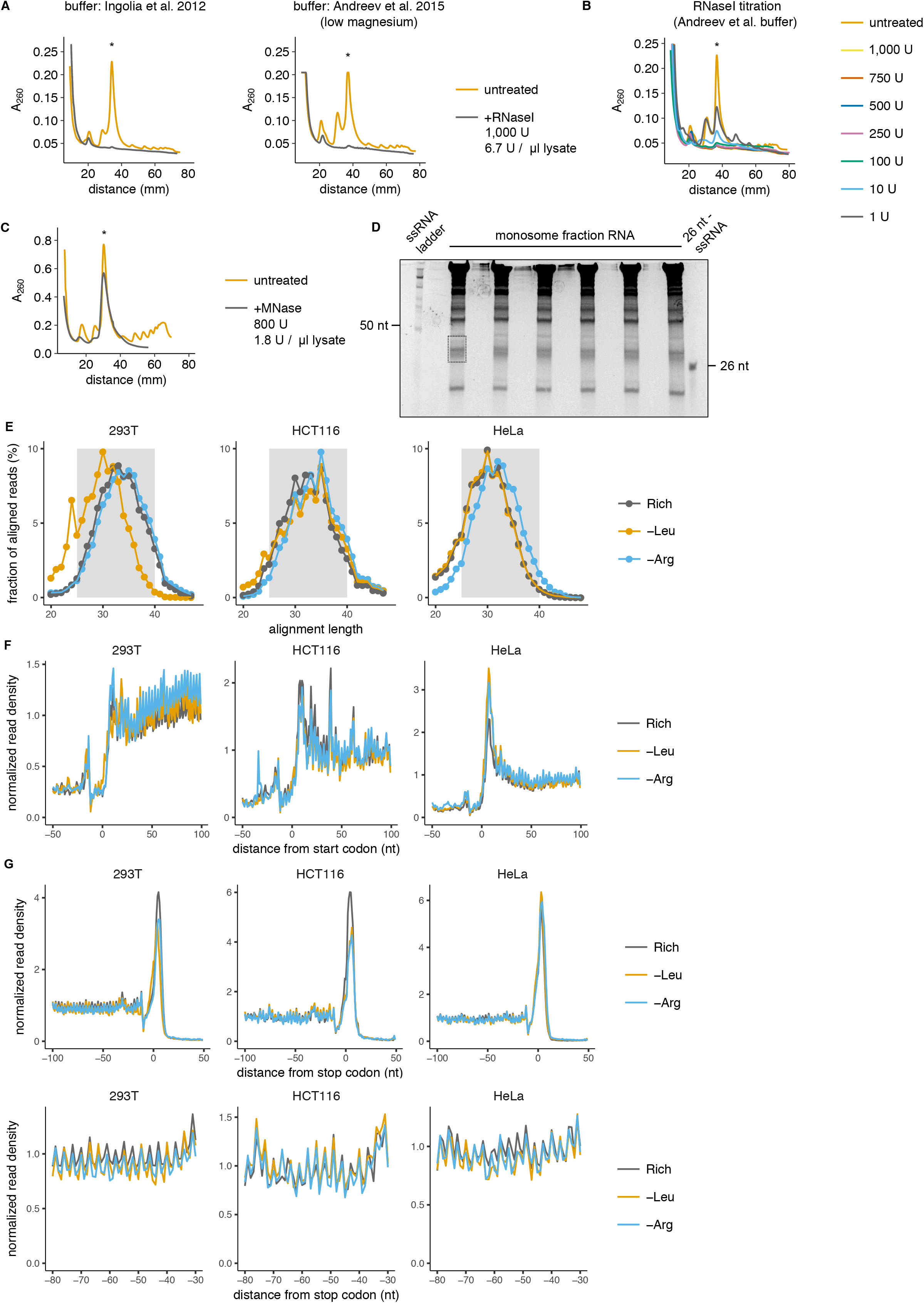
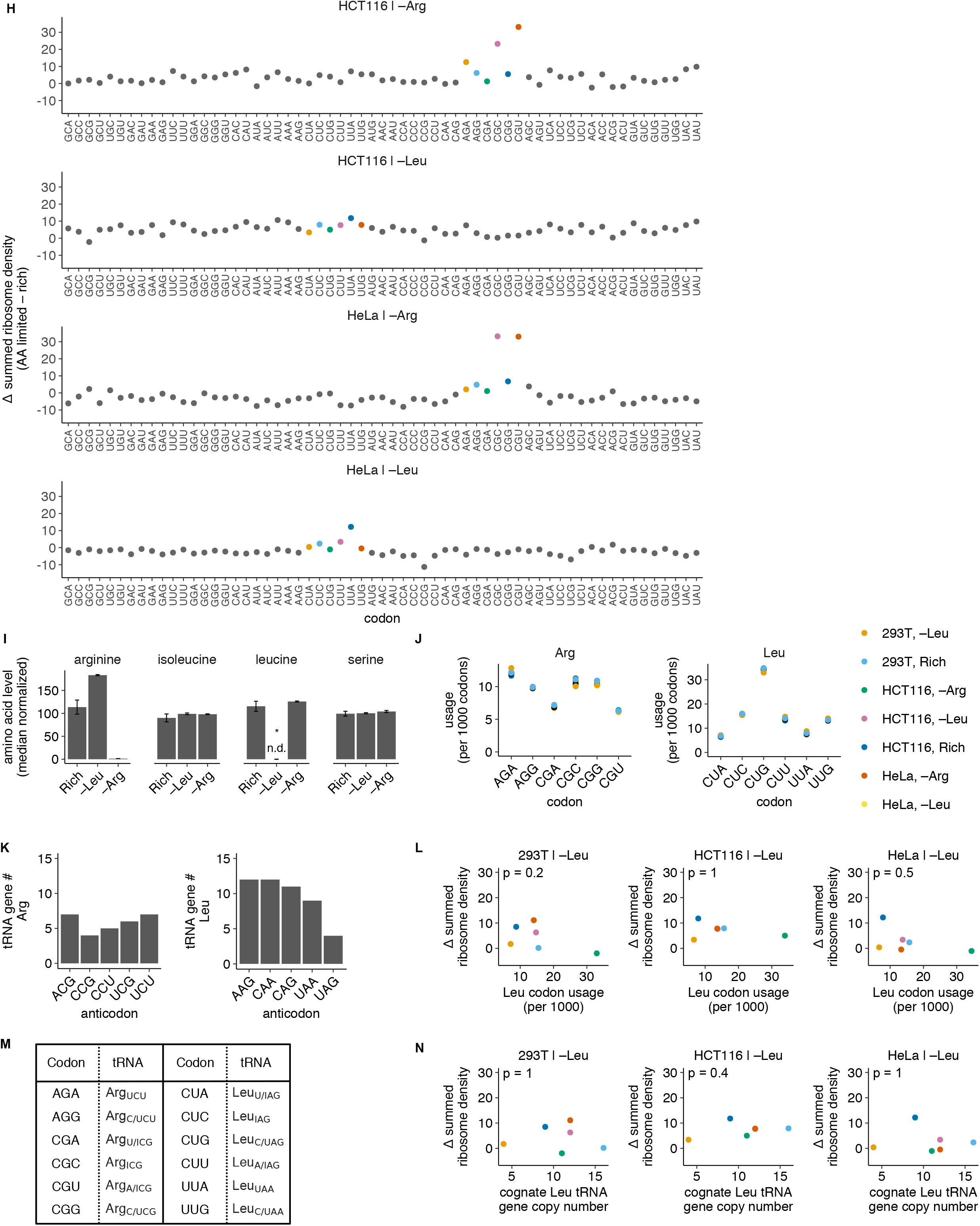
Codon-specific ribosome pausing during limitation for arginine, but not leucine. (**A-C**) HEK293T cell polysome digestion by RNaseI (A,B) or MNase (C) into monosome-bound RNA footprints assessed by sucrose density gradient fractionation. For tests with RNaseI (A,B), high and low magnesium buffers were compared. Asterisk indicates monosome fraction. (**D**) Representative 15% TBE urea size selection gel from which monosome-bound RNA footprints were extracted from the total monosome sucrose density gradient fraction for library preparation. Dashed box indicates the footprint region excised. (**E-G**) Aligned read length distribution (E) and genome-wide read density profiles around annotated start (F) and stop codons (G) used to assess library quality for ribosome profiling experiments in HEK293T, HeLa, and HCT116 cells after arginine or leucine limitation for 3 hours or growth in nutrient rich media (see Fig. 1A-C). After 3’ end trimming (see Methods), normalized read density is calculated relative to the mean footprint density for each coding sequence, and is averaged over all occurrences of the codon across detectably expressed transcripts. A region of the stop codon read density profile (G) is magnified in a second panel to clearly show three nucleotide periodicity. (**H**) Summed changes in codon-specific ribosome density for HCT116 and HeLa cells following 3 hours of arginine or leucine limitation, measured using ribosome profiling (calculated as described in Fig. 1A). (**I**) Intracellular arginine, isoleucine, leucine, and serine levels in HEK293T cells following limitation for arginine or leucine for 3 hours, relative to rich medium measured by liquid chromatography tandem mass spectrometry (LC-MS/MS). Error bars represent the standard error of the mean from three technical replicate measurements. Intracellular leucine level was below the detection limit (n.d.) upon its limitation. (**J**) Usage frequencies for Arg and Leu codons in the transcriptome in HEK293T, HCT116, and HeLa cells following 3 hours of limitation for leucine or arginine, or growth in rich conditions. (**K**) Genomic copy numbers of all Arg and Leu isoacceptor tRNAs (Chan and Lowe, 2016). (**L**) Arg and Leu codons matched with their cognate tRNA(s). Decoding by multiple tRNAs is indicated with a slash, I = inosine. (**M**) Usage frequency of Leu codons in the HEK293T, HCT116, and HeLa transcriptomes following 3 hours of leucine limitation (as shown in J) compared to the summed change in ribosome density upon leucine limitation (as shown in Fig. 1A and H). p indicates p-value of Spearman’s rank coefficient, ρ (HEK293T; ρ = –0.6, p = 0.2. HCT116; ρ = 0.03, p = 1. HeLa; ρ = –0.37, p = 0.5). (**N**) Genomic copy number of cognate tRNA for each Leu codon (as shown in K) compared to the change in ribosome density upon leucine limitation (as shown in Fig. 1A and H) (HEK293T; ρ = –0.03, p = 0.96. HCT116; ρ = 0.4, p = 0.4. HeLa; ρ = 0, p = 1).

**Supp. Fig. 2.**
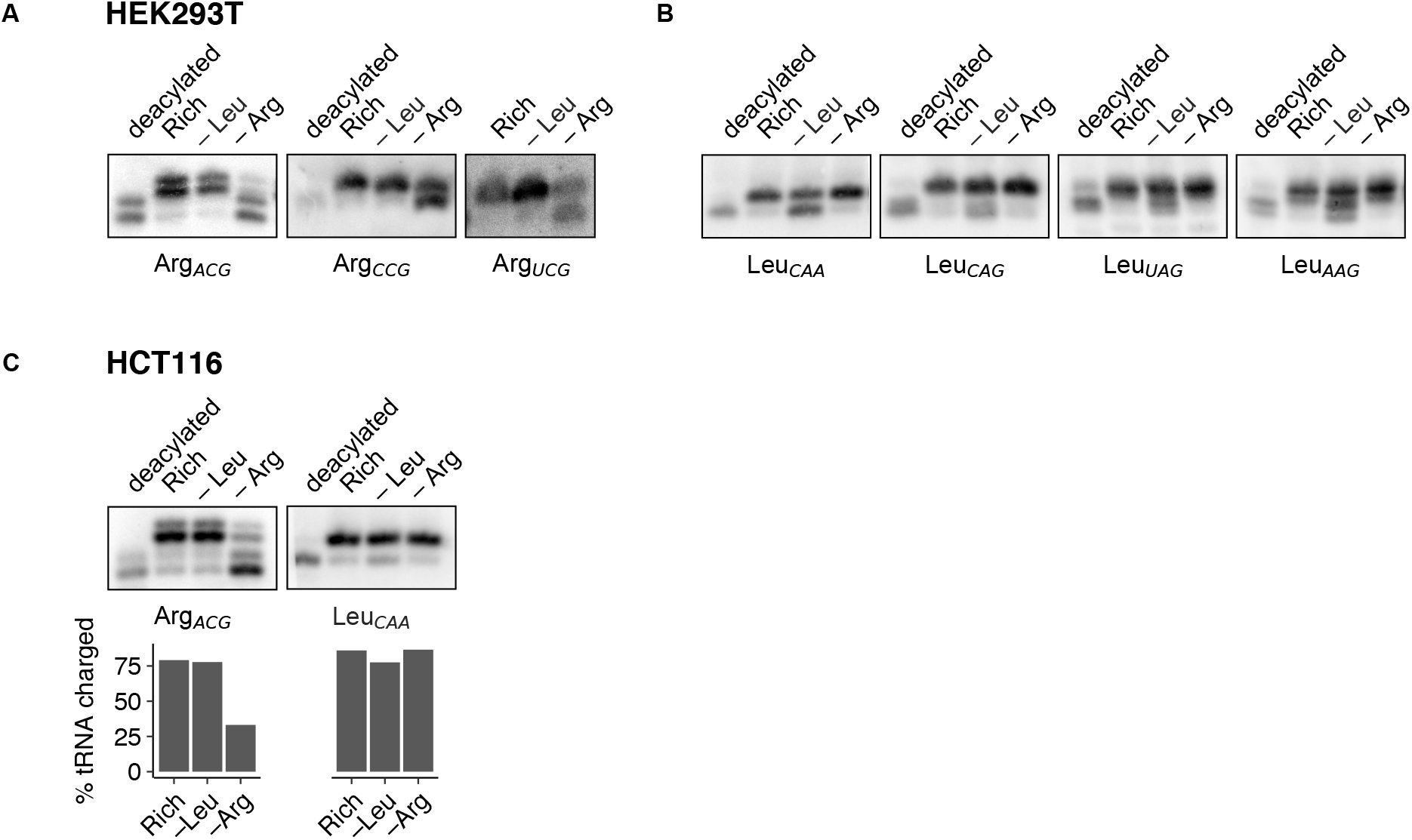
Selective loss of cognate tRNA charging during arginine limitation. (**A-C**) Representative northern blots for determination of Arg and Leu tRNA charging levels (as shown in Fig. 2) in HEK293T (A,B) cells or HCT116 cells (C) following 3 hours of limitation for arginine or leucine or growth in rich medium. A control deacylated total RNA sample is used to identify uncharged tRNA species. tRNA probe is indicated below each blot. Note that two charged and uncharged species of tRNA^Arg^_ACG_ are detected in both cell lines, likely due to covalent modification of this tRNA. Absolute charging level was calculated by dividing the intensity of the charged band(s) by the sum of all band intensities. There is a low level of cross-hybridization between the TAG and AAG probes, as we could not design unique probes for these highly homologous tRNAs (see Methods for probe design details).

**Supp. Fig. 3.**
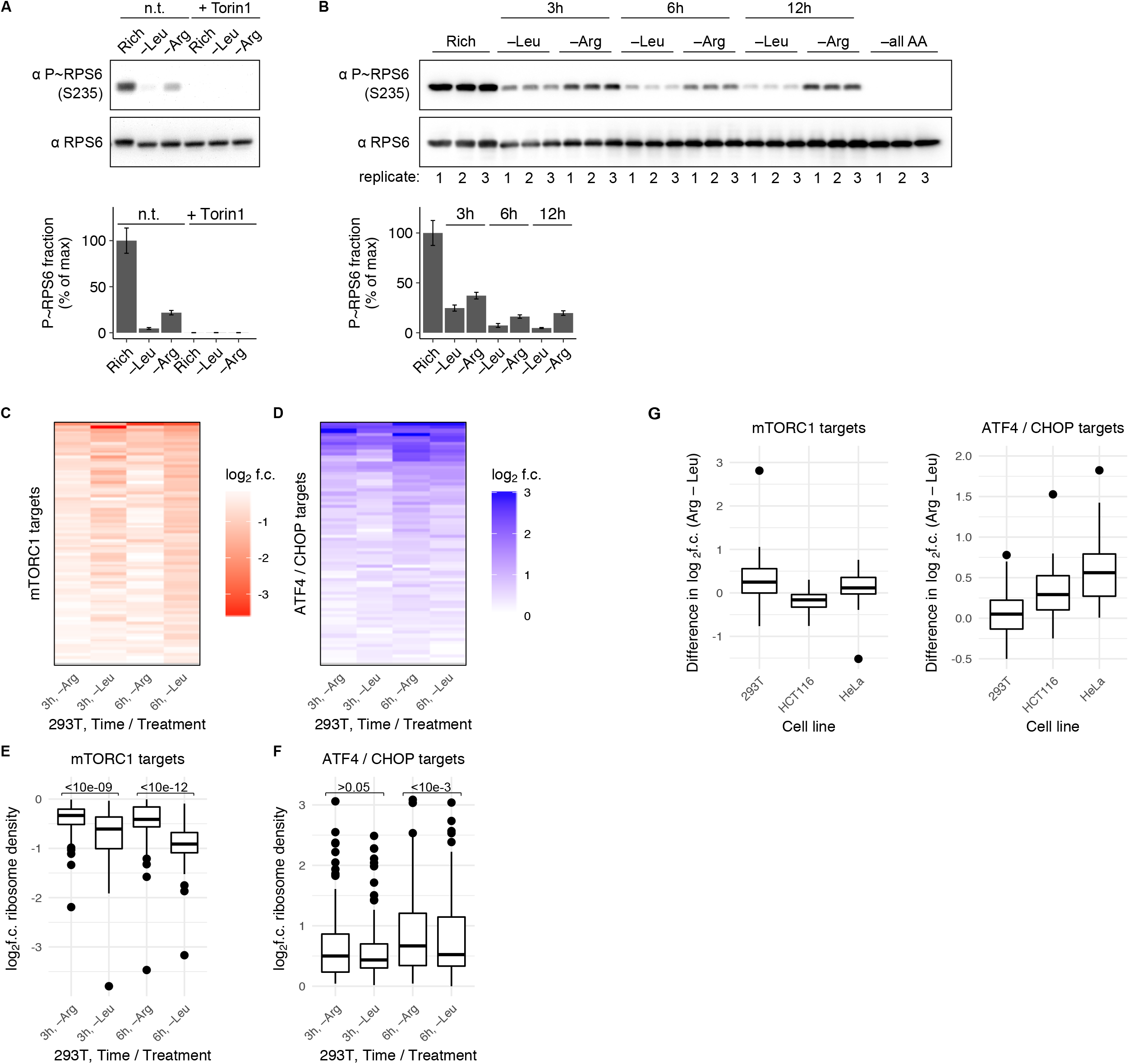
Divergent response of mTORC1 and GCN2 signaling pathways to arginine versus leucine limitation. (**A,B**) Representative western blots for phosphorylated and total levels of the S6K target, RPS6, in HEK293T cells after growth in rich medium or limitation for leucine or arginine for 3 hours in the presence or absence (n.t.) of 250 nM Torin1 (A) or after growth of three replicates in rich medium, limitation for leucine or arginine for 3, 6 or 12 hours, or limitation for all amino acids for 6 hours (B). Bar graphs show the fraction of protein that is phosphorylated in each condition, relative to rich medium; error bars represent the standard error of the mean from three technical replicate experiments. (**C,D**) Heatmap of log_2_ fold-changes (f.c.) in ribosome density for mRNA targets of translational downregulation due to mTORC1 inhibition (Hsieh et al., 2012) (C) or transcriptional or translational upregulation due to GCN2 activation (Han et al., 2013) (D) following 3 or 6 hours of arginine or leucine limitation, relative to rich medium, in HEK293T cells. Only targets with a log_2_ fold change of <0, for mTORC1, or >0, for ATF4/CHOP (the transcription factor effectors downstream of GCN2 activation), in all conditions were considered. At 3 hours, 43/73 (59%), and at 6 hours, 47/73 (64%) of mTORC1 targets had higher ribosome density upon arginine than leucine limitation. At 3 hours, 67/87 (77%), and at 6 hours, 77/87 (89%) of ATF4/CHOP targets had higher ribosome density upon arginine than leucine limitation. (**E,F**) Box plot of the log_2_ fold change for each mTORC1 (E) or GCN2 (F) target upon amino acid limitation (as shown in C,D). A two-sided Wilcoxon signed rank test with continuity correction (μ = 0) was performed (described in Fig. 3E,F legend). The resulting p-value is shown above the data for each comparison. After 3 hours versus 6 hours of limitation for arginine or leucine, the mTORC1 signaling response was 1.3- or 1.4-fold higher during arginine limitation, respectively (E) and the GCN2 signaling response was 1- or 1.1-fold higher during arginine limitation, respectively (F). (**G**) Box plot of the difference in the log_2_ fold change between each mTORC1 or GCN2 target following 3 hours of limitation for arginine versus leucine in HEK293T, HCT116, and HeLa cells.

**Supp. Fig. 4.**
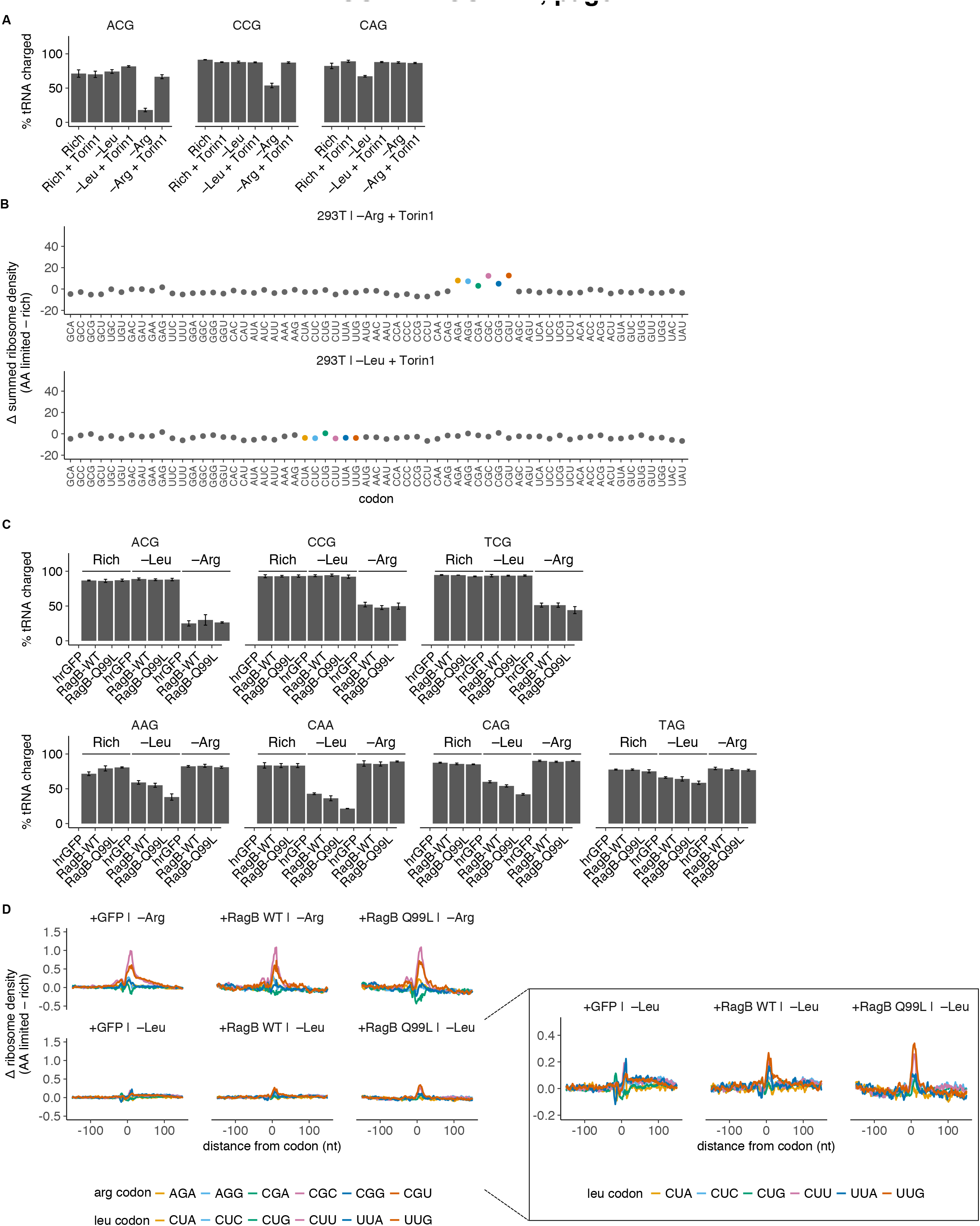
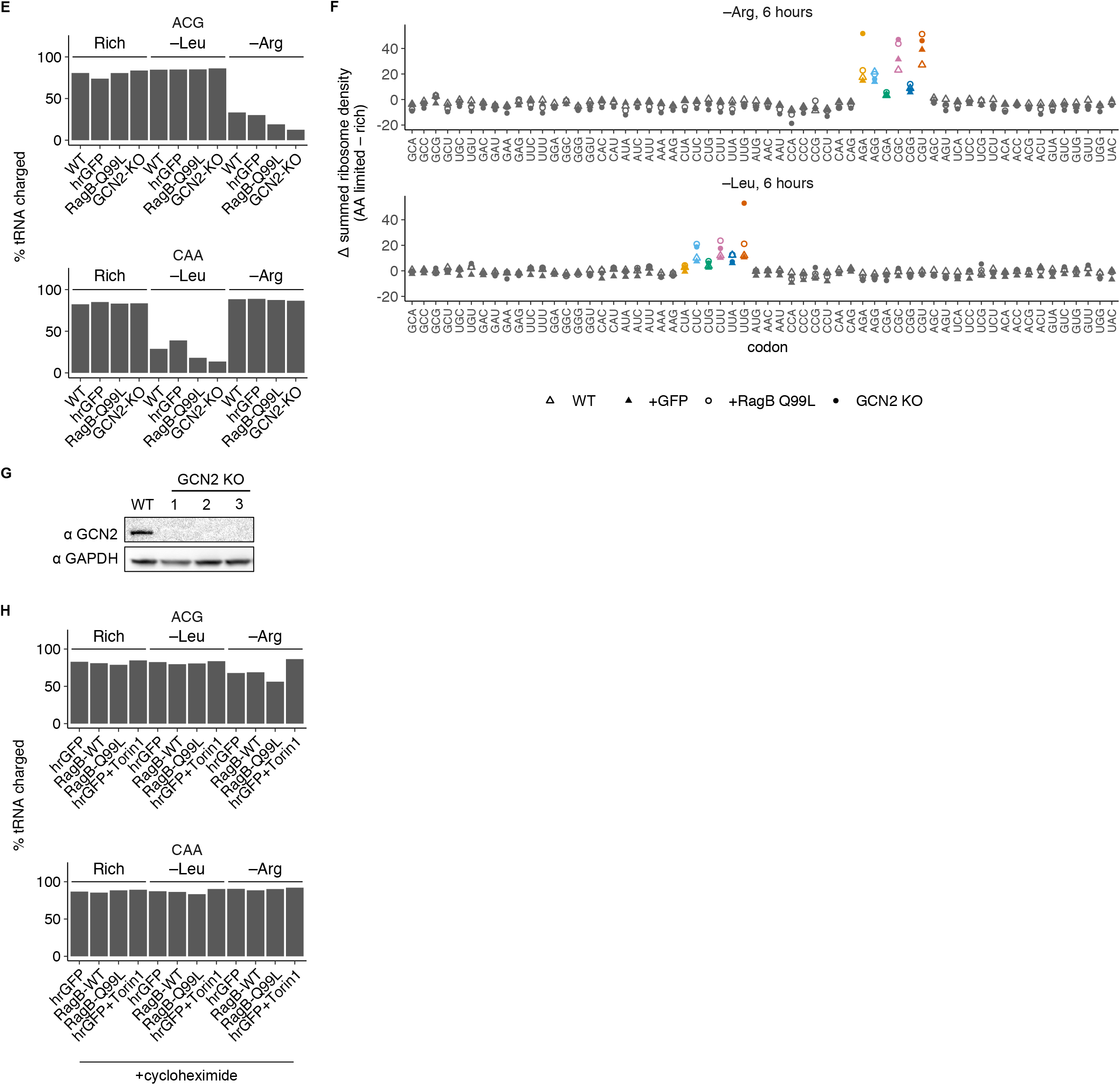
Signaling through the mTORC1 and GCN2 pathways regulates the magnitude of ribosome pausing during amino acid limitation. (**A**) tRNA charging levels for 2 Arg tRNAs and 1 Leu tRNA in HEK293T cells following 3 hours of leucine or arginine limitation or growth in rich medium, in the presence or absence of 250 nM Torin1 (calculated as described in Supp. Fig. 2A). Error bars represent the standard error of the mean from three technical replicate experiments. (**B**) Summed changes in codon-specific ribosome density for HEK293T cells expressing the fluorescent reporter protein hrGFP (Fig. 4C) following 3 hours of limitation for arginine or leucine with 250 nM Torin1, relative to rich medium. (**C**) tRNA charging levels for 3 Arg tRNAs and 4 Leu tRNAs in HEK293T cells expressing hrGFP, RagB-WT, or RagB-Q99L (as shown in Fig. 4C) following limitation for leucine or arginine for 3 hours or growth in rich medium. Error bars represent the standard error of the mean from three technical replicate experiments. (**D**) Changes in codon-specific ribosome density for the hrGFP, RagB-WT, and RagB-Q99L cell lines following limitation for leucine or arginine for 3 hours, relative to rich medium. Inset plot series shows magnified ribosome pausing around leucine codons. (**E**) tRNA charging levels for 1 Arg tRNA and 1 Leu tRNA in the WT, hrGFP, RagB-Q99L, or GCN2 KO cell lines (see Fig. 4D,E; Supp. Fig. 4G) following limitation for leucine or arginine for 3 hours or growth in rich medium. (**F**) Overlaid summed changes in codon-specific ribosome density for the WT, hrGFP, RagB-Q99L, and GCN2 KO cell lines following 6 hours of arginine or leucine limitation, relative to rich medium. (**G**) Representative western blots for GCN2 and GAPDH proteins in the WT and GCN2 KO cell lines in 3 clonal replicate GCN2 KO cell lines to verify complete protein knockout. (**H**) tRNA charging levels for 1 Arg tRNA and 1 Leu tRNA in the hrGFP cell line in the presence or absence of 250 nM Torin1, the RagB-WT and the RagB-Q99L cell line after treatment for <1 minute in ice-cold PBS with 100 µg/mL cycloheximide, following limitation for leucine or arginine for 3 hours or growth in rich medium.

**Supp. Fig. 5.**
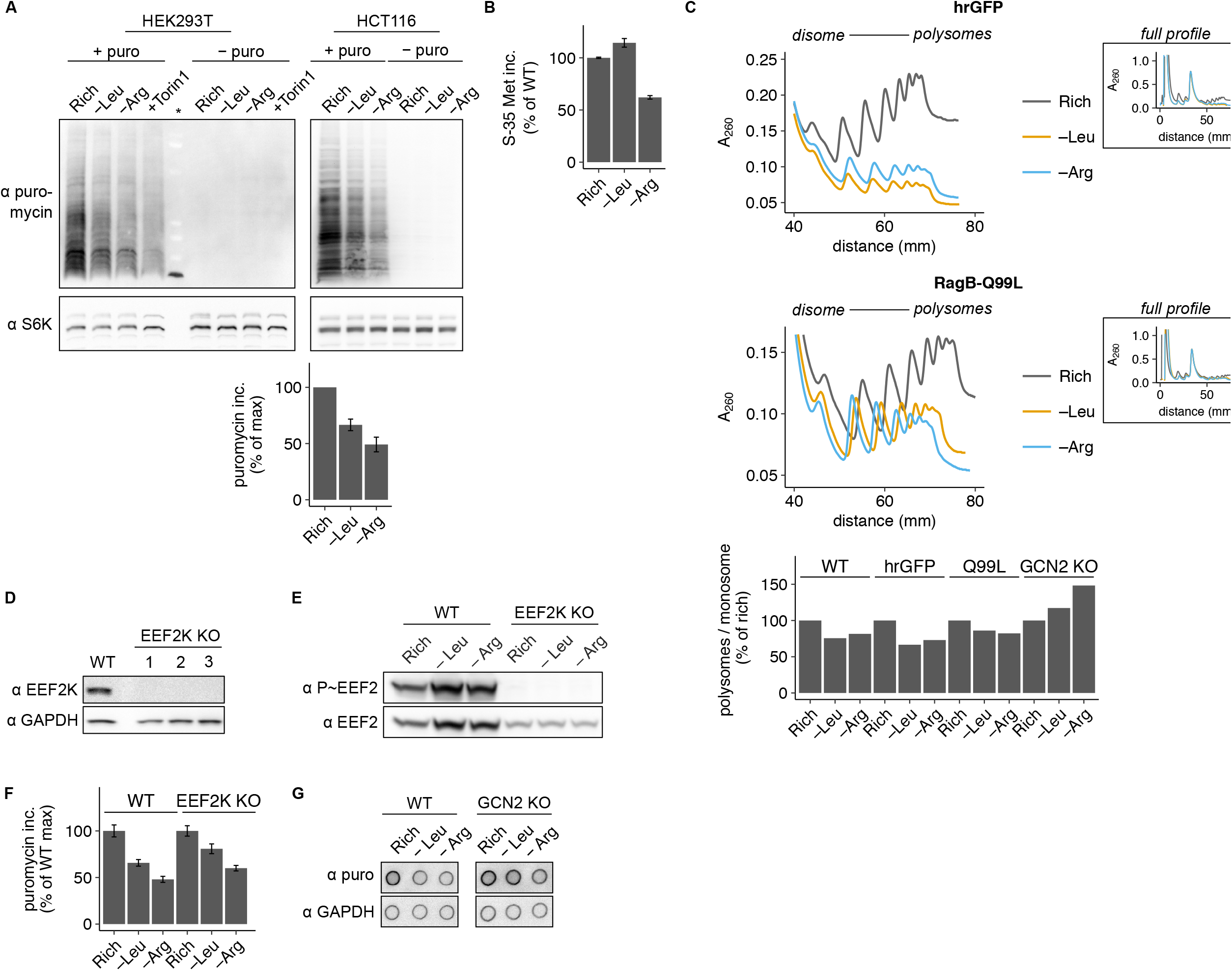
Ribosome pausing reduces global protein synthesis rate during amino acid limitation. (**A**) Representative western blots for puromycin and S6K in HEK293T (WT) and HCT116 cells given a brief pulse of 10 µg/mL puromycin (or no pulse) following 3 hours of leucine or arginine limitation, treatment with 250 nM Torin1, or growth in rich medium. To quantify global protein synthesis rate, the total puromycin signal is integrated from each lane and normalized to a western blot for total S6K protein. Bar graph shows puromycin incorporation relative to rich medium; error bars represent the standard error of the mean from three technical replicate experiments. (**B**) ^35^S-methionine incorporation into protein in the hrGFP cell line following 3 hours of leucine or arginine limitation, relative to rich medium; error bars represent the standard error of the mean from three technical replicate experiments. (**C**) Polysome profiles measured by sucrose density gradient fractionation of polysomes extracted from the hrGFP and RagB-Q99L cell lines following 6 hours of arginine or leucine limitation or growth in rich medium. The main plot shows overlaid polysome profiles from the disome (2 ribosome) peak to the end of the polysomes for all conditions, the inset plots show the entire profile. All traces were aligned with respect to the monosome peak height along the y-axis and position along the x-axis. Bar graph shows the relative area in the polysome fraction (2+ ribosomes) to the monosome fraction (1 ribosome) (see Methods for details of calculation). (**D**) Representative western blots for EEF2K and GAPDH in WT and 3 clonal replicate EEF2K KO cell lines to verify complete protein knockout. (**E**) Representative western blots for phosphorylated and total EEF2 in WT and EEF2K KO cell lines following 3 hours of growth in rich medium, leucine limitation, or arginine limitation. (**F**) Global protein synthesis rate in the WT (same data as Fig. 5D) or EEF2K KO cell lines following 3 hours of leucine or arginine limitation, relative to rich medium (calculated as described in G). Error bars represent the standard error of the mean for three technical replicate measurements. (**G**) Representative dot blots for puromycin and GAPDH in WT cells and the GCN2 KO cell line following 3 hours of leucine or arginine limitation or growth in rich medium. To quantify global protein synthesis rate, the total puromycin signal is integrated for each dot and normalized to the total GAPDH signal.

**Supp. Fig. 6.**
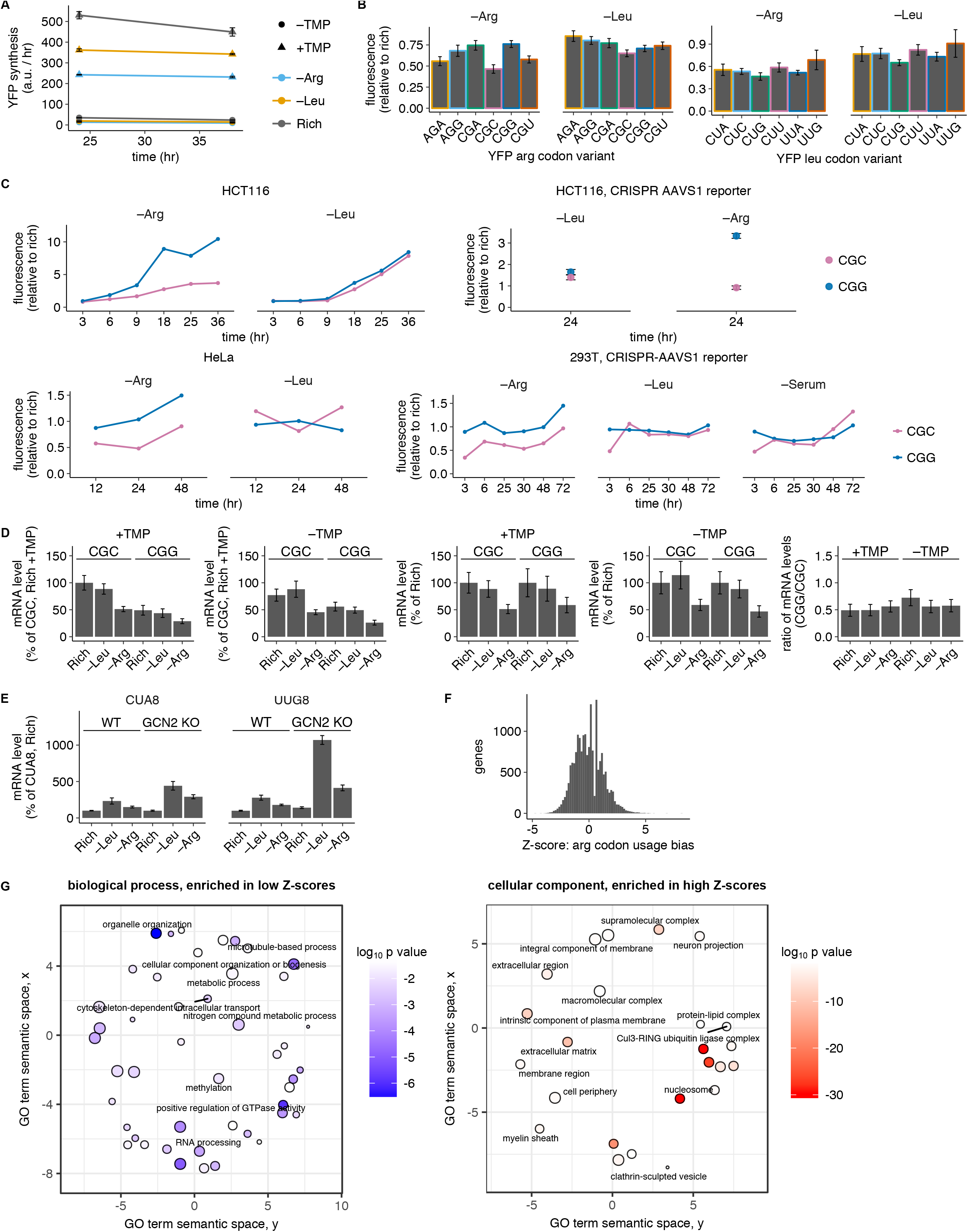
Ribosome pausing reduces protein expression from reporter mRNAs and induces premature termination of translation. (**A-C**) YFP codon variant reporter fluorescence measurements across multiple time points, cell lines, and reporter constructs. In all plots, flow cytometry was used to find the population mean YFP fluorescence from >10,000 events; error bars represent the standard error of the mean from three technical replicate experiments. (**A**) YFP fluorescence in the presence or absence of 10 µM of the reporter stabilizing ligand trimethoprim (+/-TMP) in HEK293T cells stably expressing the YFP-CGC (YFP-WT) reporter, following 24 or 38 hours of arginine or leucine limitation or growth in rich medium. (**B**) YFP fluorescence in the HEK293T cells stably expressing the arginine or leucine YFP codon variant reporters following limitation for arginine or leucine with 10 µM trimethoprim (+TMP) for 24 hours, relative to rich medium +TMP. (**C**) YFP fluorescence in the HCT116, HeLa, and HEK293T cell lines stably expressing the YFP-CGC and -CGG reporters, following limitation for arginine, leucine or serum +TMP for 12, 24, or 48 hours, relative to rich medium +TMP. Unless otherwise indicated, the reporter was introduced by lentiviral transduction (as in Fig. 6A-D, Supp. Fig. 6A,B). In the HEK293T and HCT116 cell lines, the YFP reporter constructs were also introduced by homologous recombination at the AAVS1 locus via CRISPR and contained alternative UTR and promoter elements (see Methods section under Plasmid construction for details). (**D**) YFP-CGC and -CGG reporter mRNA levels introduced at the AAVS1 locus in HEK293T cells following 24 hours of limitation for leucine or arginine in the presence or absence of TMP, relative to rich medium +TMP (see Methods section for details of calculation). From left to right, the data is displayed in a series of plots 1) without further normalization, 2) normalized to the rich condition for each YFP variant, and 3) as the ratio of the YFP-CGG variant to the YFP-CGC variant in each condition. Error bars represent the standard error of the mean for three technical replicate experiments. (**E**) CUA8 and UUG8 reporter mRNA levels in the WT and GCN2 KO cell lines following 48 hours of limitation for leucine or arginine –TMP, relative to rich medium –TMP (see Methods section for details of calculation). Error bars represent the standard error of the mean for three technical replicate experiments. (**F**) Distribution of pause-inducing arginine codons usage bias in endogenous genes (see Methods section for details). A histogram of Z-scores is shown for all coding sequences; low Z-scores represent bias against usage of pause-inducing codons to encode arginine, and high Z-scores represent bias in favor of their usage. Z-scores range from –4.7 to 8.4. (**F**) Visualization of biological process (BP) or cellular component (CC) gene ontology (GO) categories enriched in genes with bias against (left plot, BP terms enriched in lowest Z-scores) or in favor of (right plot, CC terms enriched in highest Z-scores) usage of pause-inducing codons to encode arginine using topGO (Alexa and Rahnenfuhrer, 2016; Grossmann et al., 2007) and REVIGO (Supek et al., 2011) (see Methods section for details). Each bubble represents a significantly enriched GO term; color represents log_10_ of the false-discovery rate adjusted p-value, and size scales with the number of genes associated with that term.

**Supp. Fig. 7.**
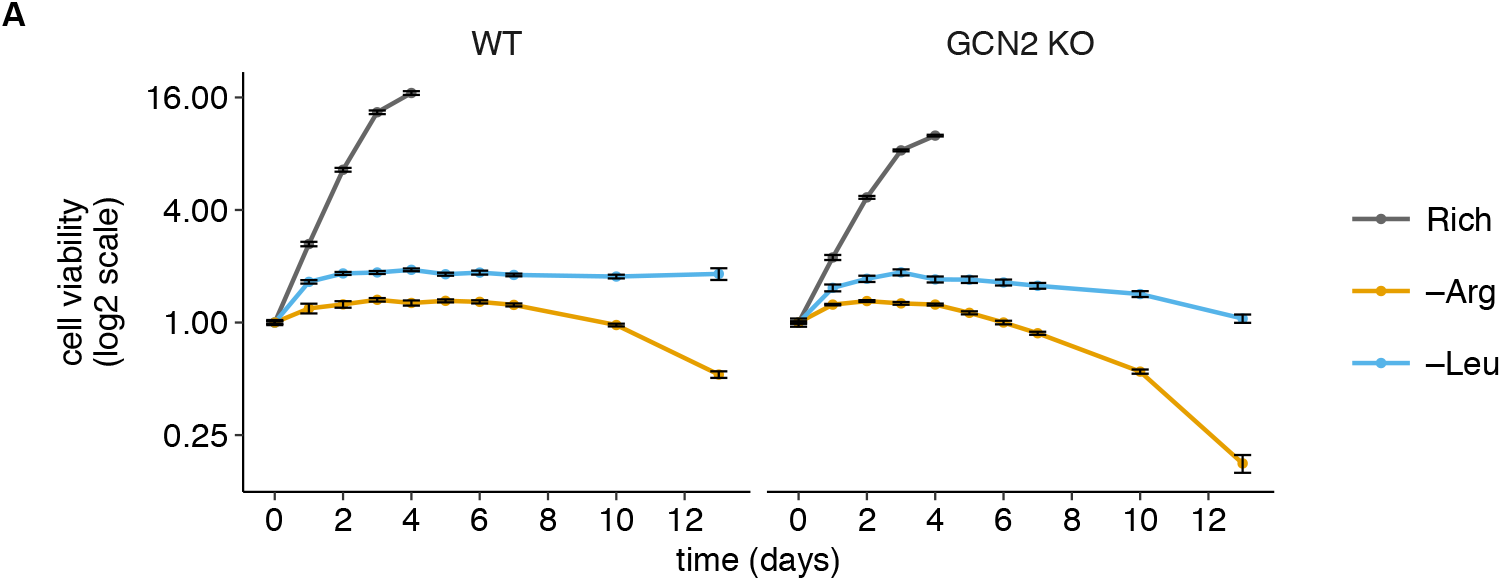
Ribosome pausing is linked to cell viability loss. (**A**) Cell viability in HEK293T cells or the GCN2 KO cell line following 1 to 13 days of arginine or leucine limitation, or growth in rich medium. The luminescence-based CellTiterGlo assay was used to find total cellular ATP content, which was normalized to the value on day 0; measurements are plotted on a log_2_ scaled y-axis. Error bars represent the standard error of the mean from five technical replicate measurements. Both cell lines reached confluency in the rich medium condition after 4 days.

## Acknowledgments

We thank B. Zid, C. Chidley, T. Pan, A. Murray, V. Denic, V. Mootha, and members of the O’Shea lab for helpful discussions. We thank C. Chidley, J. Piechura, and J. Darnell for comments on the manuscript. We thank C. Shoemaker for reagents and advice for CRISPR/Cas9 genome editing in human cell lines, and K. Han for reagents used to build the YFP codon variant protein synthesis reporters. From the FAS Division of Science, Harvard Core Facilities we thank C. Daly for high-throughput sequencing data collection and K. Chatman for LC-MS/MS data collection. The high-throughput sequencing data analysis in this paper was run on the Odyssey cluster supported by the FAS Division of Science, Research Computing Group at Harvard University. This research was supported by the National Institute of General Medical Sciences of the National Institutes of Health under award numbers, R00GM107113 and R35GM119835 (A.R.S.). EKO is an Investigator of the Howard Hughes Medical Institute. The authors declare no competing interests.

## Author Contributions

Conceptualization, A.M.D., A.R.S., and E.K.O.; Methodology, A.M.D and A.R.S.; Formal Analysis, A.M.D. and A.R.S.; Investigation, A.M.D. and A.R.S.; Writing – Original Draft, A.M.D., A.R.S., and E.K.O.; Writing – Review & Editing, A.M.D., A.R.S., and E.K.O; Funding Acquisition, A.R.S. and E.K.O.; Supervision, A.R.S. and E.K.O.

